# Structural Mechanisms of SAMD9 Autoinhibition and Pathogenic Dysregulation

**DOI:** 10.64898/2026.02.02.703423

**Authors:** Zongjun Mou, Fushun Zhang, Marisol Morales, Bibekananda Sahoo, Xinghong Dai, Yan Xiang

## Abstract

SAMD9 and SAMD9L (SAMD9/9L) are large cytosolic proteins essential for hematopoietic homeostasis and antiviral defense^1–3^. Germline gain-of-function (GoF) mutations in SAMD9/9L cause severe multisystem disorders and predisposition to leukemia, yet the mechanisms that regulate SAMD9/9L activity and how pathogenic mutations disrupt these processes, remain poorly understood. Here, we determine cryo-electron microscopy structures of human SAMD9 in multiple conformational and oligomeric states. We show that SAMD9 predominantly adopts a closed, autoinhibited conformation stabilized by a central ATP-bound nucleotide-binding oligomerization domain (NOD) and an extensive network of intramolecular interactions. Recurrent patient-derived GoF mutations localize to and destabilize these intramolecular interfaces, and restoring the disrupted interactions through compensatory mutations reinstates autoinhibition. We further identify low-abundance asymmetric SAMD9 dimers in which one protomer undergoes large conformational changes and establishes intermolecular interactions that are essential for SAMD9 activation. Together, these findings define the structural basis of SAMD9 autoinhibition and explain how human GoF mutations subvert this regulatory mechanism to drive disease.

## INTRODUCTION

Sterile Alpha Motif Domain-containing 9 (SAMD9) and its paralog SAMD9-like protein (SAMD9L) (collectively SAMD9/9L) are large cytosolic proteins encoded in tandem on human chromosome 7 and play essential roles in cell growth control, hematopoietic homeostasis, and antiviral defense^1–3^. Heterozygous mutations in either gene cause a spectrum of human disorders, including MIRAGE syndrome^4,5^, ataxia-pancytopenia^6,7^, immunodeficiency^8,9^, and systemic autoinflammatory diseases^9,10^, and collectively account for a large fraction of inherited bone marrow failure syndromes^8,11–13^. Pathogenic alleles are predominantly missense gain-of-function (GoF) mutations that enhance the intrinsic growth-suppressive and translation-inhibitory activities of SAMD9/9L^4,8,14,15^. These mutations act in an autosomal dominant manner, and affected individuals frequently acquire monosomy 7 as a somatic “rescue” to escape growth restriction^5,7,8^. This adaptive process paradoxically increases the risk of developing myelodysplastic syndromes and leukemia^16^.

SAMD9/9L are also recognized as antiviral factors against an expanding range of viruses, including poxviruses^17–19^, HIV-1^20^, rotaviruses^21^, and flaviviruses^22^. The evolutionary significance of this function is underscored by the fact that poxviruses encode dedicated antagonists of SAMD9/9L; for example, vaccinia virus (VACV) lacking the viral inhibitors K1 and C7 fails to replicate in mammalian cells due to abortive protein synthesis^18,19,23^.

Bioinformatic and functional studies suggest that SAMD9/9L belong to the Signal Transduction ATPases with Numerous Domains (STAND) superfamily of signaling proteins^14,24^, which are defined by a central nucleotide-binding oligomerization domain (NOD) flanked by variable N-terminal effectors and C-terminal regulatory modules^25^. This superfamily includes key regulators of cell death and innate immunity, such as Apaf-1, NOD-like receptors (NLRs), and plant disease-resistance (R) proteins. Canonical STAND proteins typically exist in an ADP-bound, inactive state and undergo ADP-to-ATP exchange, NOD reorganization, and oligomerization upon sensing pathogen- or stress-associated signals^26,27^.

Recent work identified the SAMD9/9L effector as a tRNA endoribonuclease (tRNase) located within their N-terminal region^14,28^. The tRNase is activated during poxvirus infection, selectively depleting tRNA^Phe^ and inducing translational arrest^28^. Importantly, patient-derived GoF mutations render the tRNase constitutively active even in the absence of a viral trigger, resulting in chronic translational repression and cellular dysfunction^28^. However, the mechanisms by which this enzymatic activity is regulated within full-length SAMD9, and how it is activated by pathogens or by diverse GoF mutations, have remained unknown.

Here, we combine cryo-electron microscopy (cryo-EM) with structure-guided mutagenesis and cellular assays to define the molecular basis of SAMD9 regulation. We establish SAMD9 as a *bona fide* STAND protein, yet one that uses ATP binding and lineage-specific structural elements to maintain the autoinhibited state. By capturing low-abundance asymmetric dimers, we also reveal partially open conformations that represent activation intermediates. Our analyses reveal that disruption of key intramolecular interactions is a major pathogenic mechanism underlying GoF mutations and that specific intermolecular interactions are essential for SAMD9 activation.

## RESULTS

### Cryo-EM analysis uncovers multiple conformational and oligomeric states of SAMD9

Purification of full-length human SAMD9 yielded variable amounts of protein due to substantial retention on chromatography columns (Fig. S1A). The N-terminal ∼150 residues contain a SAM domain followed by a low-complexity segment, and deletion of this region had only a minor effect on SAMD9 antiviral activity (Fig. S1B-C). We therefore expressed a truncated construct beginning at residue 156 (SAMD9^156–1589^) in HEK293-F cells. HaloTag affinity purification followed by TEV protease cleavage and size-exclusion chromatography yielded highly pure, monodispersed protein that displayed tRNA^Phe^ cleavage activity (Fig. S1D-G).

Cryo-EM analysis of SAMD9^156–1589^ revealed that most particles corresponded to monomers (38%) or symmetric homodimers with two-fold rotational (C2) symmetry (49%). Exhaustive 3-dimentional (3D) classification further identified two low-abundance asymmetric dimers, which we term “shell” dimer (5%) and “wing” dimer (8%) based on their overall shapes (Fig. 1A, Fig. S2, Table S1). In each asymmetric dimer, one protomer was structurally indistinguishable from the monomer, whereas the second protomer adopted markedly different conformations, with some domains poorly resolved or absent due to increased conformational flexibility (Fig. 1A, S2D).

**Figure 1.**
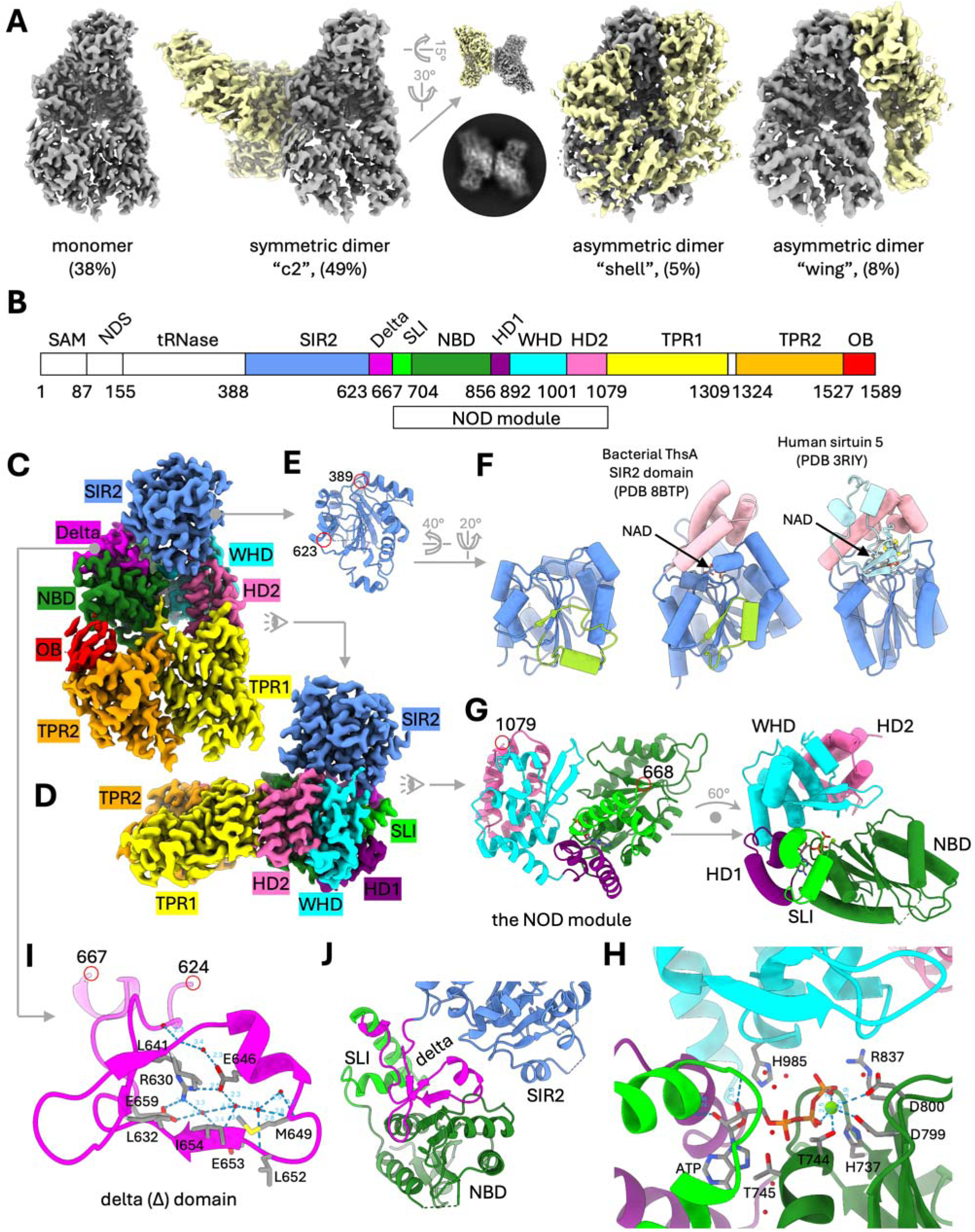
Cryo-EM reconstruction and domain architecture of human SAMD9. (A) Cryo-EM density maps of the four major SAMD9 particle classes identified by three-dimensional classification. A representative two-dimensional class average of C2 dimers is also shown. (B) Domain organization of SAMD9 from N-to C-terminus. (C, D) Cryo-EM structure of the SAMD9 monomer in different views, with domains colored as in panel B. (E, F) Structure of the SIR2-like domain. In (F), the SIR2-like domain is compared with that of the prokaryotic anti-phage protein ThsA^32^ and human sirtuin 5^33^. (G, H) Structure of the NOD module. In (H), a close-up view of the ATP-binding pocket highlights key nucleotide-interacting residues. (I, J) Structure of the Δ domain. The Δ domain mediates interactions between the SLI-NBD region and the SIR2-like domain.

### Overall architecture of the SAMD9 monomer

Based on sequence analysis and the cryo-EM structures, we defined the SAMD9 domain architecture as follows (Fig. 1B): sterile alpha motif (SAM) domain, N-terminal disordered segment (NDS), tRNase domain, Silent information regulator 2 (SIR2)-like domain, Δ domain, NOD module, and a C-terminal region consisting of tetratricopeptide repeat 1 (TPR1), a short flexible linker, TPR2, and an oligonucleotide/oligosaccharide-binding (OB) domain. In all cryo-EM reconstructions, the first resolved residue corresponds to the N-terminus of the SIR2 domain, indicating that the tRNase and SIR2 domains are flexibly linked. The SAMD9 monomer adopts an L-shaped conformation reminiscent of a chaise longue, with the SIR2 domain forming an upright “backrest” and the subsequent domains curling around to create an elongated, horizontal “seat” (Fig. 1C-D).

The SIR2-like domain exhibits a three-layered α/β Rossmann fold corresponding to the large domain of canonical SIR2 proteins^29^, which are NADases containing an additional smaller domain that forms the NAD-binding lid (Fig. 1E-F). Accordingly, the SAMD9 SIR2-like domain is unlikely to bind NAD. Structurally, it more closely resembles recently described prokaryotic SIR2-like proteins involved in diverse antiphage defense systems than eukaryotic sirtuins^30,31^. Specifically, both SAMD9 SIR2 and prokaryotic SIR2-like proteins lack the three-stranded zinc-binding element characteristic of eukaryotic sirtuins and instead share an insertion that includes a β-strand augmenting the core β-sheet (shown in yellow green in Fig. 1F).

Positioned directly underneath the SIR2 domain, the NOD module contains all four canonical domains: the nucleotide-binding domain (NBD), helical domain 1 (HD1), winged-helix domain (WHD), and helical domain 2 (HD2). The NBD and WHD resemble their counterparts in Apaf-1 and NLRP3, whereas HD1 and HD2 are reduced in size, consisting of only two and three α-helices, respectively, rather than the four and five or more helices typically found in other STAND NODs (Fig. 1G, S3).

Immediately N-terminal to the NOD module lies the Δ domain, named for its triangular shape (Fig. 1I-J). The domain is rigidified by an extensive hydrogen-bonding network formed by the sidechains of R630, E646, and E659, together with several ordered water molecules and backbone atoms (Fig. 1I). The Δ domain mediates interactions between the NBD and SIR2 domains, positioning SIR2 above the cleft between the NBD-HD1 and WHD-HD2 submodules (Fig. 1C, J).

C-terminal to the NOD module are two TPR arrays of α-helical bundles separated by a flexible linker, followed by an OB domain with a five-stranded β-barrel. Together, the TPR-OB region curls back toward the NBD, forming the body of the “seat” (Fig. 1C, 2A).

**Figure 2.**
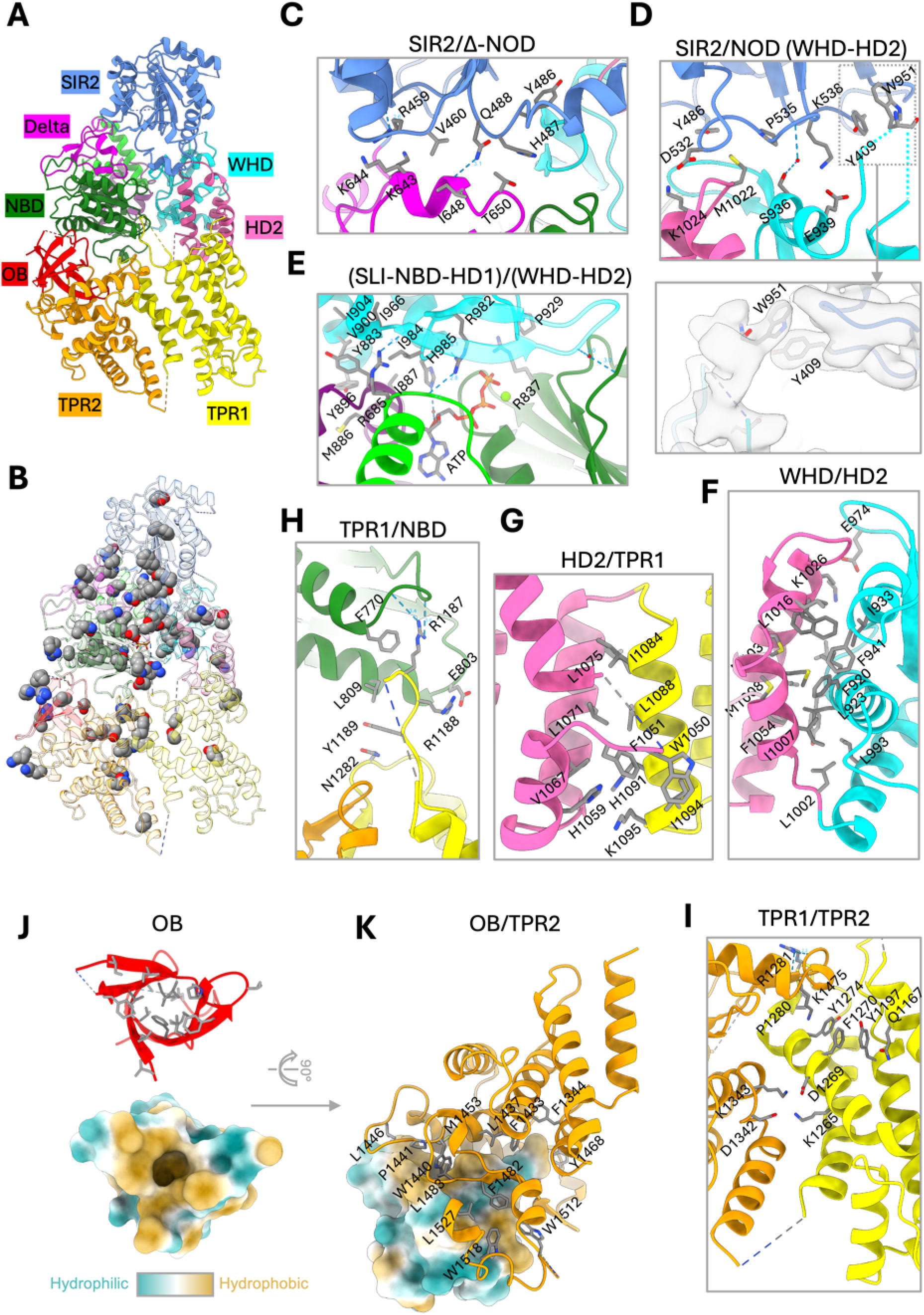
The closed SAMD9 conformation is reinforced by extensive intramolecular interactions. (A) The atomic model of SAMD9 in the autoinhibited state, with individual domains colored as in Fig. 1B. (B) Mapping of known patient-derived GoF mutations onto the SAMD9 atomic model (shown as spheres). Many GoF mutations cluster at intramolecular domain-domain interfaces. (C-I) Close-up views of representative intramolecular interfaces, highlighting key stabilizing interactions between the indicated domains. Inset in (D) shows well-resolved sidechain density for W951 stacking against Y409, despite poor backbone density for the loop containing W951. (J) Structure of the OB domain highlighting the hydrophobic core of the β-barrel. The lower panel shows surface representation colored by hydrophobicity, illustrating a deep hydrophobic pocket at the base of the β-barrel. (K) Hydrophobic interactions between the OB domain and the TPR2 domain. The sidechain of W1440 from TPR2 inserts into the hydrophobic pocket of the OB domain.

### SAMD9 monomer and C2 dimer adopt an ATP-bound closed conformation

Clear densities for an ATP and a Mg^2+^ ion are observed at the center of the NOD module in both the monomer and C2 dimer (Fig. 1H, S4H). Because ATP was not added during purification, SAMD9 must stably bind endogenous cellular ATP and hydrolyzes it very slowly, if at all. Strikingly, despite ATP occupancy, the NOD adopts a closed conformation resembling the ADP-bound, inactive states of Apaf-1 and NLRP3, with the WHD positioned against the bound nucleotide, rather than the ATP-bound, open conformations associated with activation (Fig. S3).

ATP engagement by the NOD broadly follows the canonical configuration^34^, involving the Walker A and Walker B motifs, the Sensor-1 residue, and a WHD histidine (Fig. 1H), all of which are conserved across the mammalian SAMD9/9L family (Supplemental file 1). However, the Walker A motif is noncanonical: the invariant lysine of the GKT/S consensus is replaced by glycine (^742^GGT^744^). Instead, the sidechain of an upstream H737 occupies the spatial position normally taken by this lysine and likely fulfills a similar functional role. T744 within the Walker A motif contacts the Mg^2+^ ion, which in turn coordinates the β- and γ-phosphates of ATP. Two consecutive acidic residues in the Walker B motif, D799 and D800, also contact the Mg^2+^ ion, while the Sensor-1 residue R837 is positioned near the γ-phosphate of ATP (Fig. 1H).

Supporting the importance of these interactions, mutations T744A, D799A, D800A, or R837A resulted in loss of function (LoF) and markedly reduced SAMD9 antiviral activity, whereas the T745A substitution had a smaller effect (Fig. S4A-C). This contrasts with Walker B mutations in NLRP3, which cause spontaneous activation^35^. Attempts to purify the T744A and D800A variants resulted in substantially reduced protein purity and yield (Fig. S4D), suggesting that ATP binding is important for SAMD9 folding or stability. In contrast, we successfully purified and determined the cryo-EM structure of the R837A mutant (Fig. S4E-H). ATP remains bound in this structure, consistent with observations from Sensor-1 mutants in other ATPases, which typically retain nucleotide binding but display impaired hydrolysis^34^. Notably, the R837A dataset consisted entirely of C2 dimers (Fig. S4F), lacking both monomers and the asymmetric dimers.

In canonical STAND proteins, a WHD histidine (e.g., H438 in Apaf-1, H522 in NLRP3, and H443 in NLRC4) stabilizes the autoinhibited state by contacting the β-phosphate of ADP and tethering WHD and NBD domains together (Fig. S3B-C); substitution of this residue with Leu in NLRC4 causes spontaneous activation^36^. This histidine in inactive NLRP3 would have clashed with ATP^37^, rationalizing the large WHD rotation and NOD opening that accompany ADP-to-ATP exchange. In SAMD9 NOD, however, an N-terminal helix-coil-helix element (residues 667-704), which we term the SAMD9 Lineage Insert (SLI), wedges between the NBD and WHD, slightly displacing the WHD and creating a wider NOD cleft than is observed in inactive Apaf-1 or NLRP3 (Fig. S3A-C), providing sufficient space to accommodate ATP. The corresponding WHD histidine, H985, interacts with ATP via coordinated water molecules (Fig. 1H), and the H985A mutation resulted in a GoF phenotype (Fig. S5), indicating that H985 plays a regulatory role analogous to that of the conserved WHD histidine in other STAND proteins.

The C2 dimer consists of two identical protomers that retain the closed conformation and associate through a flat interface along the side of the SLI-NBD-HD1 region. Dimerization is mediated by P704/P704 and R863/Y701 stacking interactions and three pairs of salt bridges (Fig. S6), interlocking SLI-NBD-HD1 and potentially reinforcing the ATP-bound closed state.

Together, these results indicate that ATP binding is essential for SAMD9 stability and autoinhibition, whereas stable ATP engagement without proper γ-phosphate sensing may lock SAMD9 in the autoinhibited state.

### The closed SAMD9 conformation is reinforced by extensive intramolecular interactions

To date, approximately 95 distinct pathogenic SAMD9 variants have been identified in patients^1^. To investigate their pathogenic mechanisms, we mapped these mutations onto the monomeric SAMD9 structure (Fig. 2A-B). Most of the mutations are predicted to perturb intramolecular interactions or disrupt local conformations (Table S2), thereby destabilizing the closed state or facilitating the conformational changes required for activation.

Nearly half of the variants are located to the SIR2-Δ-NOD region. SIR2 interacts with NOD indirectly through the Δ domain and directly with the WHD-HD2 region. The Δ domain plays a critical role in mediating SIR2/NBD interactions. A β-strand at the base of the Δ domain augments the central NBD β-sheet (Fig. 1J), while a cluster of hydrophobic residues reinforce tight packing against the NBD. A short helix at the shoulder of the Δ domain mediates interactions with the SIR2 domain (Fig. 1J, 2C). Notably, GoF variants R459Q and K643E map to this interface.

SIR2/(WHD-HD2) interactions involve a helix (residues 934-942), a partially flexible loop (residues 943-955), and β-turn (residues 972-978) within the WHD, as well as a HD2 coil (residues 1019-1023). W951 on the WHD loop stacks against Y409 in SIR2 (Fig. 2D), and a recurrent GoF variant E974K maps to the WHD β-turn.

Within the NOD, the NBD-HD1 and WHD-HD2 submodules face each other and are bridged by ATP, ordered water molecules, and a network of long sidechains, most prominently R685, R982, and H985 (Fig. 2E). Notably, R685Q and R982H/C are GoF mutations, and we show that the H985A also results in a GoF phenotype (Fig. S5).

HD2 serves as a spacer that couples the NOD to the C-terminal region. The extensive hydrophobic interactions between HD2 and WHD (Fig. 2F) suggest that these domains act together as a rigid body, transmitting conformational changes of the WHD to the C-terminal region, as observed in other STAND proteins^37^.

The HD2/TPR1 interface is stabilized by both hydrophobic and polar interactions, including two facing histidines (H1059 and H1091), suggesting a potential sensitivity to pH (Fig. 2G). A partially flexible HD2 loop (residues 1036-1048) further reinforces this interface and harbors a recently identified GoF variant, G1048R^38^, which may disrupt loop binding by reducing backbone flexibility.

TPR1 and TPR2 form a scissor-like configuration hinged via two short β-strands. This architecture is stabilized by stacking interactions and salt bridges, with GoF mutations (P1280L, Y1274C) perturbing these contacts (Fig. 2I). In addition, a partially flexible TPR1 loop (residues 1173-1192) interacts with NBD and harbors multiple GoF mutations (Fig. 2H).

The OB domain is tightly anchored to TPR2 via hydrophobic interactions at both the base and periphery of its β-barrel (Fig. 2J-K). Although OB seems spatially separated from the NBD in the whole-dataset reconstruction, local classification revealed that an OB loop (residues 1557-1569) frequently contacts the NBD near R824 (Fig. S7), a recurrent GoF site (R824Q) (Table S2).

### Disruption of key intramolecular interfaces represents a major pathogenic mechanism underlying SAMD9 GoF mutations

Among the pathogenic variants, five recurrent GoF mutations have been identified in more than four independent cases^1^, and all are predicted to disrupt intramolecular domain-domain interactions on the monomeric structure (Table S2). One such recurrent mutation, E974K, is predicted to destabilize the SIR2/(WHD-HD2) interface (Fig. 3A). E974 is located on the SIR2-interacting WHD β-turn, forming a charge-charge interaction with K1026 in HD2. We therefore hypothesized that restoring a similar charge-charge interaction would be sufficient to stabilize the WHD β-turn harboring the E974K mutation and suppress spontaneous activation. To test this idea, we introduced the K1026E substitution into both the wild-type (WT) and E974K background. Strikingly, K1026E alone exhibited a strong GoF effect comparable to E974K, whereas the E974K&K1026E double mutant restored autoinhibition (Fig. 3B-E).

**Figure 3.**
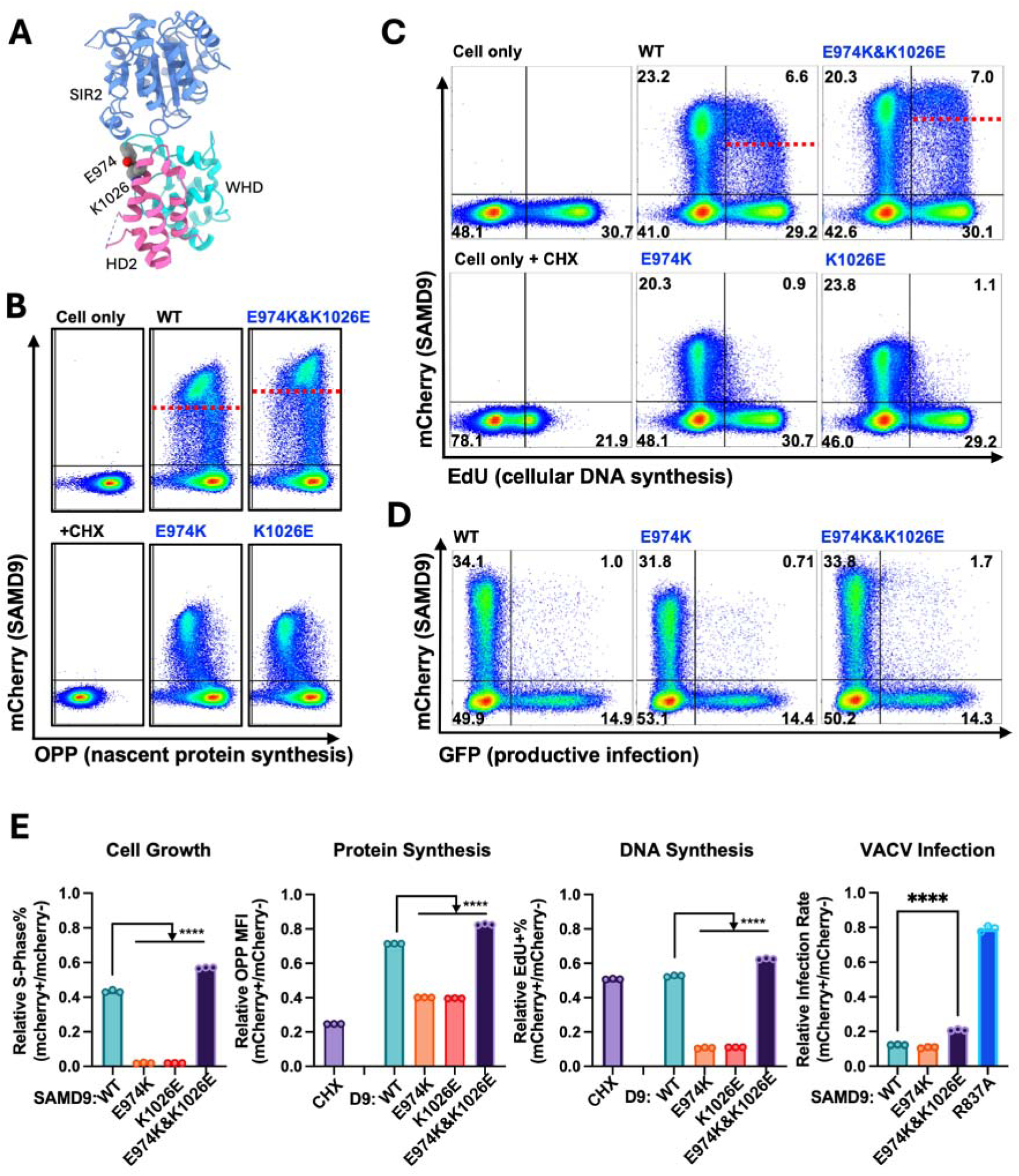
Restoration of the SIR2/(WHD-HD2) interface suppresses pathological activation by the GoF E974K mutation. (A) Structural view of the SIR2/(WHD-HD2) interface highlighting the electrostatic interaction between E974 in the WHD (cyan) and K1026 in HD2 (pink). (B-D) HEK293T cells were transfected with mCherry-SAMD9/9L fusion constructs. (B) Cells were labeled with O-propargyl-puromycin (OPP) for 30 min to measure nascent protein synthesis. Representative flow-cytometry plots show OPP incorporation versus mCherry expression. Untransfected cells in the absence or presence of cycloheximide (CHX) are shown as controls. The red dashed line indicates the mCherry-SAMD9 expression threshold above which inhibition of protein synthesis is observed. (C) Cells were labeled with EdU (2 h pulse) to assess DNA synthesis. Representative flow-cytometry plots show EdU incorporation versus mCherry expression. The red dashed line indicates the mCherry-SAMD9 expression threshold above which inhibition of DNA synthesis is observed. (D) Cells were infected with a SAMD9-susceptible VACV mutant, vK1^-^C7^-^/GFP□. Representative flow-cytometry plots show virus-encoded GFP expression versus mCherry expression, with quadrant percentages indicated. (E) Relative S-phase fractions, nascent protein synthesis, cellular DNA synthesis, and infection rates in SAMD9-expressing versus non-transfected cells within the same sample were quantified from the flow-cytometry data in panels B-D and Fig. S8A. Statistical significance was assessed by pairwise Student’s *t* tests (****, *P* < 0.0001).

While WT SAMD9 only modestly reduced cell progression into S phase, both E974K and K1026E strongly blocked S-phase entry (Fig. 3E and S8A). Moreover, whereas WT SAMD9 inhibited cellular protein synthesis and DNA replication only when its expression in cells exceeded a threshold level (indicated by the red dashed line in Fig. 3B-C), both E974K and K1026E single mutants exerted these inhibitory effects in a dose-dependent manner across the full range of expression levels. By contrast, WT and GoF mutants inhibited VACV replication with similar efficiency even at low expression levels (Fig. 3D, S8B), reflecting SAMD9 activation by viral infection. Compared to the WT, the E974K&K1026E double mutant showed a slightly reduced ability to inhibit cellular protein synthesis, DNA replication, cell-cycle progression, and VACV infection (Fig. 3B, C, E). Together, these results demonstrate that the E974/K1026 electrostatic interaction is essential for maintaining SAMD9 autoinhibition and that disruption of this single contact is sufficient to trigger pathological activation.

The SLI region appears to play a central role in stabilizing the closed NOD conformation by tethering the NBD and WHD together, and this short region is particularly enriched in GoF mutations, including R685Q. In the WT protein, R685 forms two hydrogen bonds (∼3 Å) with the backbone carbonyls of residues 965 and 967, stabilizing the NBD/WHD interface (Fig. 4C left panel). To test whether strengthening this interface could suppress the GoF phenotype, we introduced G686C and I968C into the R685Q background, aiming to engineer a disulfide bond to covalently tether the NBD and WHD. The resulting R685Q&G686C&I968C triple mutant indeed displayed a LoF phenotype and lacked antiviral activity (Fig. 4A-B).

**Figure 4.**
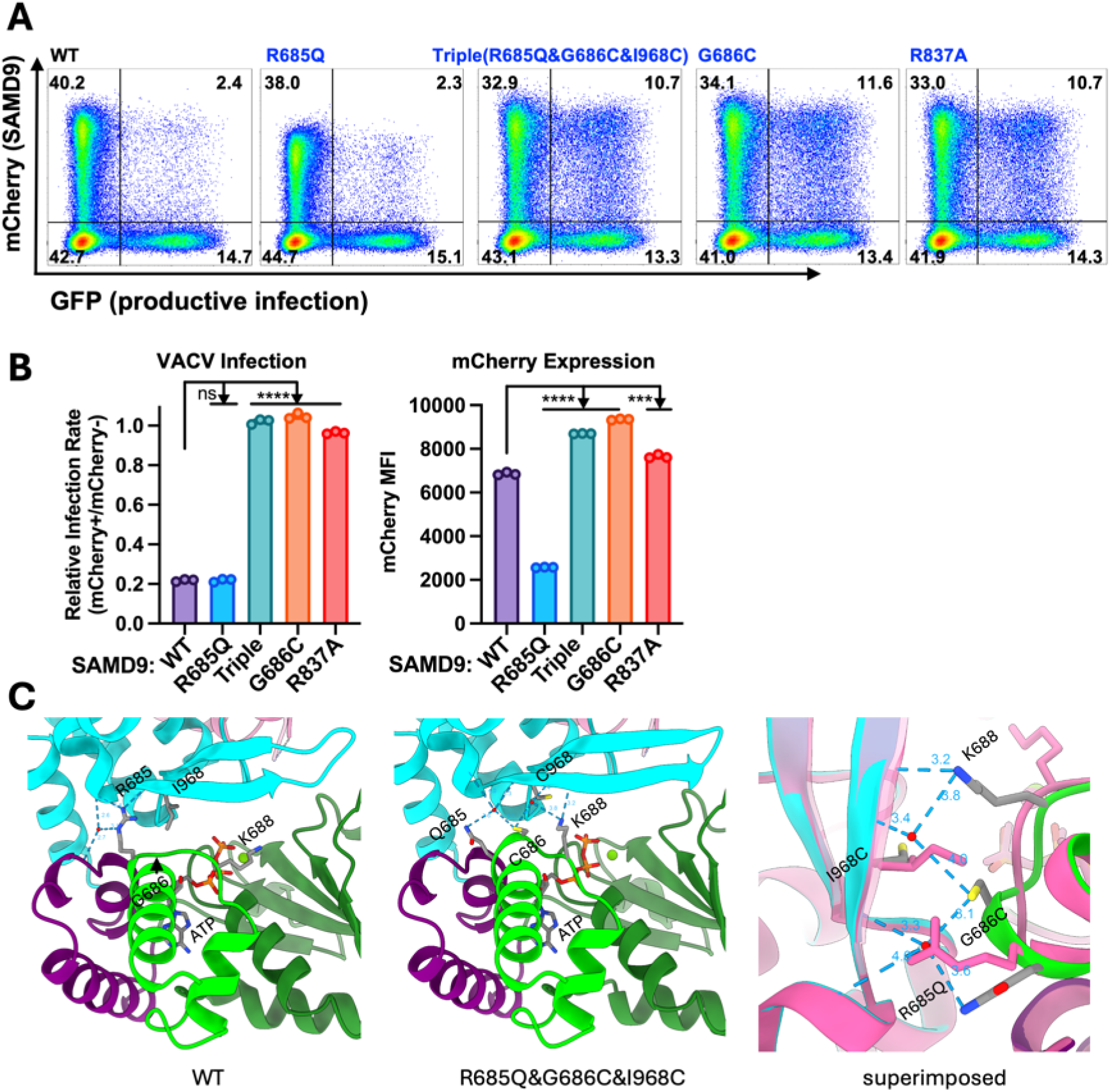
Reinforcing the NBD/WHD interface suppresses SAMD9 activation by the GoF R685Q mutation. (A) HEK293T cells were transfected with mCherry-SAMD9 fusion constructs and infected with vK1^-^C7^-^/GFP□ VACV. Representative flow-cytometry plots show virus-encoded GFP expression versus mCherry expression, with quadrant percentages indicated. (B) Quantification of VACV infection rates and mCherry-SAMD9 mean fluorescence intensity (MFI) in cells expressing the indicated SAMD9 constructs. Infection rates were normalized to mCherry-negative cells within the same sample. The mCherry-SAMD9 MFI reflects the ability of the variants to inhibit cellular protein synthesis. (C) Left, structural view of the NBD/WHD interface in WT SAMD9 highlighting R685 and neighboring residues that stabilize interdomain packing through hydrogen bonding and water-mediated interactions. Center, cryo-EM structure of the R685Q&G686C&I968C triple mutant illustrating a local rearrangement induced by the G686C substitution, which engages with two ordered water molecules and repositions K688 to form a network of compensatory hydrogen bonds with the WHD backbone. Right, overlay comparison of the WT (pink) and mutant (green and cyan) structures.

Cryo-EM analysis of the triple mutant revealed both the structural impact of the R685Q GoF mutation and the rescue mechanism mediated by the G686C mutation (Fig. 4C). The overall structure of the triple mutant closely resembled that of WT SAMD9 (Fig. S9A, C-D); however, the R685Q substitution abolished the native hydrogen bonds to the WHD backbone due to the shorter and less flexible glutamine sidechain, thereby weakening NBD/WHD contacts. Contrary to our original design, the G686C and I968C substitutions did not form a disulfide bond. Instead, the G686C sidechain engaged two ordered water molecules, one of which recruited the sidechain of K688. Together, these groups formed multiple hydrogen bonds with the WHD backbone, effectively re-stabilizing and even strengthening the NBD/WHD interface (Fig. 4C middle and right panels). Consistent with this structural observation, the G686C substitution alone was sufficient to abolish SAMD9 antiviral activity (Fig. 4A-B). Notably, asymmetric dimers were nearly absent in the cryo-EM dataset of the triple mutant (Fig. S9C).

Collectively, these results demonstrate that modulating domain-domain contacts through single amino acid substitutions can either promote or suppress GoF effects. Disruption of key intramolecular interfaces therefore represents a major pathogenic mechanism underlying SAMD9 GoF mutations.

### Asymmetric dimers capture large conformational changes in SIR2 and C-terminal region

The low-abundance asymmetric dimers of WT SAMD9 became absent in LoF R837A and R685Q&G686C&I968C mutants, suggesting that these dimers may represent activation intermediates. In the asymmetric dimers, one protomer remains in the closed conformation, whereas the other undergoes a dramatic rearrangement in which the SIR2 domain disengages from the NOD module and instead forms extensive intermolecular contacts with the SIR2 domain of the opposing protomer (Fig. 5A, E). The OB domain also swings outward and reorients to engage either its own SIR2 domain or that of the opposing protomer, giving rise to the wing and shell dimers, respectively (Fig. 5A, E). Notably, despite the markedly different overall architectures of the two dimer types, the local structures of the SIR2/SIR2 and OB/SIR2 moieties are superposable between the two types, indicating highly specific interactions. The comparable abundance of the wing and shell dimer types further suggests that the OB stochastically associates with the SIR2 from either promoter.

**Figure 5.**
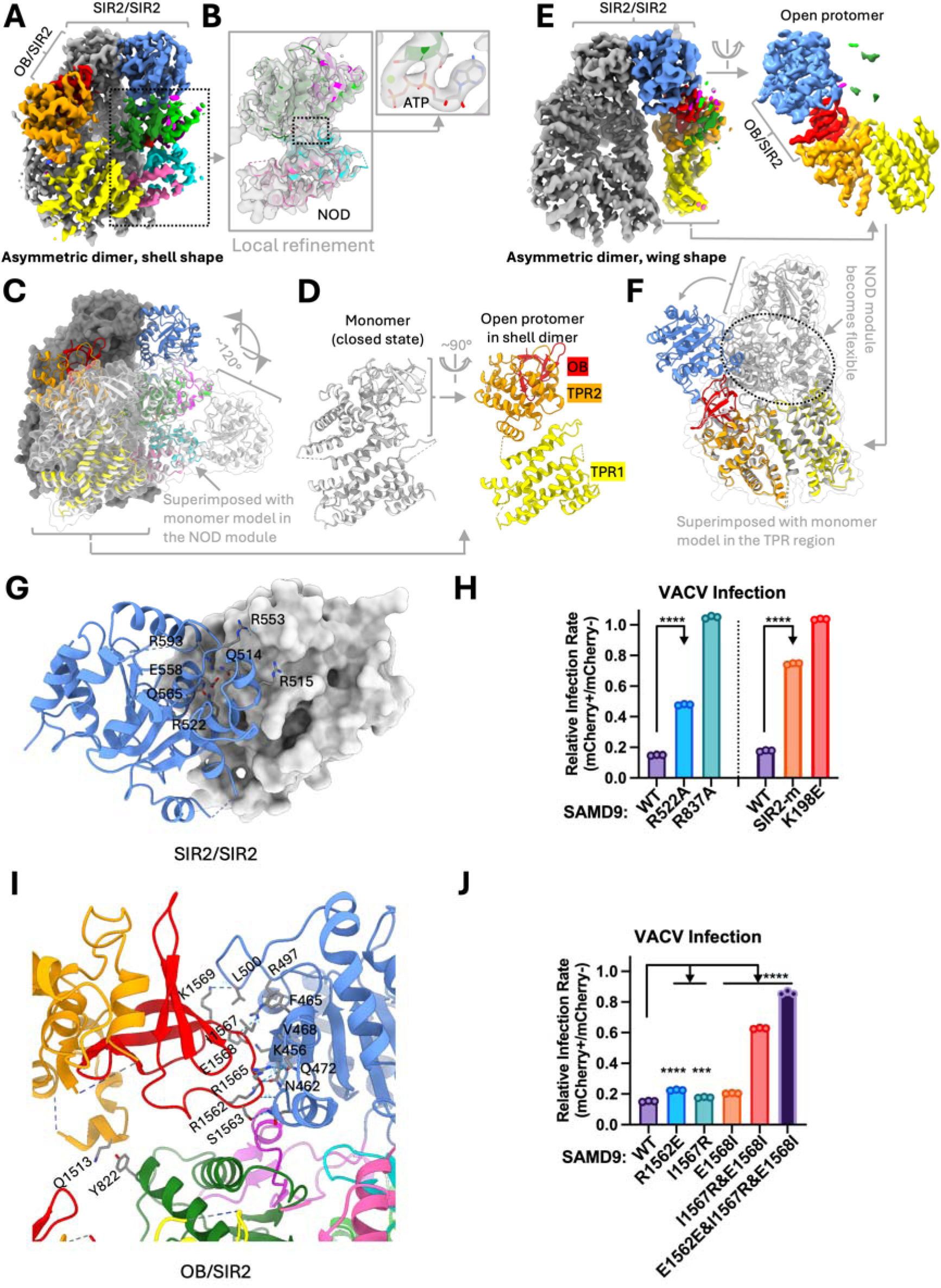
SAMD9 conformational landscapes revealed by asymmetric dimer structures and the requirement for specific intermolecular interactions in SAMD9 activation. (A) Cryo-EM density map of the asymmetric “shell” dimer, with domains of the open protomer color-coded as in Figure 1B. (B) Local refinement of the NOD module of the open protomer reveals no structural change in the NOD and that ATP remains bound. (C, D) Conformational rearrangements in the open protomer of the shell dimer. The closed protomer is shown in gray surface, and the open protomer is shown in color-coded ribbon and superimposed with a closed-state model (white) aligned at the NOD module. The SIR2 domain undergoes a ∼120° rotation to form the intermolecular SIR2/SIR2 interface, while the TPR2-OB region swings ∼90° to contact the SIR2 of the closed protomer. (E) Cryo-EM density map of the asymmetric “wing” dimer, with domains of the open protomer color-coded as in Figure 1B. (F) Conformational rearrangements in the open protomer of the wing dimer. The open protomer model is aligned to a closed-state protomer at the TPR region, revealing rotation of the SIR2 domain to engage with its own TPR-OB domains. The NOD module of the open protomer was unresolved due to increased flexibility. (G) Structure of the SIR2/SIR2 dimer interface observed in both types of asymmetric dimers. One of the SIR2 domains is shown in surface mode to illustrate the flat dimerization interface. (H) Mutations disrupting the SIR2/SIR2 dimerization interface markedly reduce SAMD9 antiviral activity. Relative VACV infection rates (% of GFP□ cells) in SAMD9-expressing versus non-transfected cells were quantified. *SIR2-m* denotes a combined mutation of six surface residues (Q514S, R515S, R522S, R553S, Q556S, and R593S). R837A and K198E are verified LoF mutants serving as negative controls. (I) Structure of the OB/SIR2 interface. Only intermolecular OB/SIR2 interactions of the shell dimer are shown here; the intramolecular OB/SIR2 interactions in the open protomer of the wing dimer are nearly identical. (J) Mutations disrupting the OB/SIR2 interaction strongly impair SAMD9 antiviral activity. Relative VACV infection rates in SAMD9-expressing versus non-transfected cells were quantified. Statistical significance in (H) and (J) was determined using pairwise Student’s *t* tests (****, *P* < 0.0001; ***, *P* < 0.001).

For simplicity, we refer to the two promoters within the asymmetric dimers as the closed promoter and the open promoter, reflecting the more open conformation of the latter. Comparison of these two protomers illustrates the dramatic structural movements that SAMD9 can undergo. In the shell dimer, the open protomer is fully resolved despite slightly increased flexibility of the NOD and TPR1 domains (Fig. 5A). Local refinement of the NOD module followed by rigid-body fitting with closed-state NOD model revealed that ATP remained bound and that there was no NOD domain reorganization (Fig. 5B) as observed in other activated STAND proteins^37,39^. When the open protomer is superposed with the closed monomer in the NOD module, the SIR2 domain rotates by ∼120° at the junction between SIR2 and NOD, and the TPR2-OB module undergoes an ∼90° rotation at the TPR1/TPR2 hinge, resulting in a complete dissociation from TPR1 (Fig. 5C-D). In the wing dimer, the open protomer is only partially resolved, with the NOD module becoming completely flexible while the TPR1-TPR2-OB remain rigidly associated and interact with its own SIR2 via OB (Fig. 5E). When superposed with the closed monomer in the TPR1-TPR2-OB region, the SIR2 domain of the open protomer also exhibits an ∼120° rotation (Fig. 5F). Together, these asymmetric dimers demonstrate that both the SIR2 and TPR2-OB regions are capable of large-scale rigid-body movements and formation of new intermolecular interactions.

### SIR2/SIR2 and OB/SIR2 interactions are critical for SAMD9 activation

SIR2/SIR2 dimerization is a defining feature of both types of asymmetric dimers. Cryo-EM analysis of a maltose-binding protein (MBP) fusion with SAMD9^623–1589^ via a flexible linker revealed monomeric and C2 dimeric assemblies of SAMD9^623–1589^, with the MBP moiety unresolved due to conformational flexibility (Fig. S9G-H). Importantly, no asymmetric dimers were observed, indicating that the SIR2 domain is indispensable for asymmetric dimerization.

The SIR2 dimerization interface is relatively flat and enriched in polar sidechains. Unusually, the interactions are dominated by planar sidechain stacking, most prominently the reciprocal stacking of the guanidinium groups of R522 and R593 from opposing protomers. In addition, the amide groups of Q514 and Q565 sidechains stack against backbone peptide bonds in the opposing subunit (Fig. 5G).

Consistent with the functional importance of SIR2 dimerization, the R522A mutation significantly impaired SAMD9 antiviral activity (Fig. 5H, S10A), whereas substitution of six surface residues with serine (Q514S, R515S, R522S, R553S, Q556S, and R593S; SIR2-m) nearly abolished activity (Fig. 5H, S10B). Cryo-EM analysis of the SIR2-m mutant revealed preservation of the overall WT structure but exclusive formation of monomers and C2 dimers, with no detectable asymmetric dimers (Fig. S9I-J), indicating selective disruption of SIR2/SIR2 interactions. In a subset of SIR2-m particles, the SIR2 domain was also flexible and not resolved (Fig. S9J), suggesting that SIR2 can dynamically engage and disengage from the NOD in equilibrium. The absence of such particles in the WT dataset further suggests that SIR2/SIR2 dimerization stabilizes this intermediate as the open protomer in the asymmetric dimer, thereby shifting the equilibrium toward activation.

The OB/SIR2 interaction is mediated by the same OB loop (residues 1557-1569) that can dynamically engage the NBD in the SAMD9 monomer. When bound to SIR2, this loop adopts a conformation similar to its NBD-disengaged state but distinct from its NBD-bound state (Fig. S7D-E), suggesting that release from the NBD primes the loop for SIR2 engagement. Residues 1562-1569 form extensive contacts with the SIR2 surface, including hydrogen bonds, salt bridges, and hydrophobic interactions (Fig. 5I). Mutation of three residues (R1562E&I1567R&E1568I) abolished antiviral activity, whereas the double mutant I1567R&E1568I or the single substitution R1562E caused partial impairment (Fig. 5J, S10C). Cryo-EM analysis of the triple mutant showed an intact overall structure but exclusive formation of C2 dimers, with no detectable monomers or asymmetric dimers (Fig. S9E-F).

Together, these results demonstrate that both SIR2/SIR2 and OB/SIR2 interactions are required for generating asymmetric dimers and for SAMD9 activities.

## Discussion

Our studies provide two key insights into the structural principles governing SAMD9 autoinhibition and activation. First, we determine the structure of SAMD9 in its autoinhibited state and establish that it is a *bona fide* STAND protein whose autoinhibition is enforced by lineage-specific structural features. Structure-guided mutagenesis further demonstrates that disruption of key intramolecular contacts constitutes a major pathogenic mechanism underlying patient-derived GoF mutations. Second, we capture partially open conformations within asymmetric dimers, revealing large-scale conformational changes accessible to specific SAMD9 domains. We further demonstrate that the intermolecular interfaces within asymmetric dimers are required for activation, establishing these dimers as *bona fide* activation intermediates and enabling us to propose a mechanistic pathway for SAMD9 activation.

### SAMD9 autoinhibition and divergence from canonical STAND proteins

SAMD9 autoinhibition departs from the classical STAND paradigm in two fundamental respects. First, the SAMD9 NOD is stabilized in an ATP-bound, rather than ADP-bound, inactive state. Second, the closed SAMD9 conformation is further reinforced by structural elements unique to the SAMD9 lineage, including SLI, the Δ domain, and a prokaryotic SIR2-like domain. Together, these elements assemble into an extensive network of intramolecular interfaces that clamp SAMD9 in a closed configuration.

Consistent with this architecture, a large fraction of patient-derived GoF mutations map to the SIR2-Δ-NOD regions, indicating that these structural elements are particularly important for maintaining autoinhibition and/or facilitating the conformational changes required for activation. This mutational distribution contrasts with that of NLRP3, in which GoF mutations are largely confined to the NOD domain^40^.

Structure-guided mutagenesis confirms that interdomain interactions are critical for autoinhibition. At the SIR2/(WHD-HD2) interface, a conserved electrostatic interaction between E974 and K1026 anchors the closed state. Both residues are highly conserved across mammalian SAMD9/9L orthologs (Supplemental file 1), indicating an evolutionarily conserved autoinhibitory mechanism. We show that K1026E is a GoF mutation comparable in severity to the recurrent E974K mutation. Notably, K1026E has not been identified in patient cohorts, underscoring the predictive power of the SAMD9 structures for identifying potential GoF alleles.

Direct structural analysis of GoF mutants has so far been challenging because they aggregate during purification. Nevertheless, by combining the GoF mutation R685Q with stabilizing substitutions that confer LoF phenotypes, we were able to determine cryo-EM structures that reveal structural perturbations caused by the R685Q mutation, even though our current approach would not capture any subsequent large-scale conformational changes induced by the GoF mutations.

### Asymmetric dimers as early activation intermediates

Our second major insight comes from structural snapshots of early activation intermediates and the identification of specific intermolecular interactions as key drivers of SAMD9 activation. Cryo-EM analysis captures asymmetric SAMD9 dimers that reveal large-scale rigid-body movements of the SIR2 and TPR2-OB, while the core NOD module remains largely intact and ATP-bound at least in the shell dimer. This differs markedly from fully activated oligomeric STAND assemblies, in which ADP-to-ATP exchange and extensive NOD domain rearrangement occur. In activated Apaf-1 and NLRs (including NLRP3 and NLRC4), the WHD-HD2 submodule rotates by ∼80-180° at the junction between HD1 and WHD, opening the NOD domains for oligomerization^37,39,41,42^.

Multiple lines of evidence indicate that the asymmetric dimers represent *bona fide* activation intermediates. First, asymmetric dimer formation disrupts SIR2/NOD interactions, consistent with the observation that mutations affecting SIR2 and NOD interactions are GoF. Second, TPR1 and TPR2 undergo large conformational rearrangements in the shell dimer, and the hinge region that couples these domains in the autoinhibited state is highly enriched in GoF mutations. Third, asymmetric dimers are markedly reduced or absent in LoF mutants. Finally, targeted disruption of the SIR2/SIR2 or OB/SIR2 interactions abolishes asymmetric dimer formation and results in loss of function.

### A switch-like model for SAMD9 activation

Integrating our findings with established STAND activation paradigms, we propose a switch-like activation model for SAMD9. In the resting state, SAMD9 undergoes transient, partial opening that involves conformational changes in the SIR2 and C-terminal domains. When another SAMD9 molecule is in proximity, intermolecular SIR2/SIR2 and OB/SIR2 interactions can stabilize the open protomer, giving rise to the low-abundance asymmetric dimers observed in our cryo-EM dataset. The closed protomer within such an asymmetric dimer can itself undergo transient opening and be stabilized by an additional SAMD9 molecule, thereby propagating conformational changes and promoting oligomerization. These intermolecular interactions may further drive rearrangements within the NOD module, exposing the NBD surface for oligomerization. Such assemblies would bring multiple tRNase domains into proximity, enabling coordinated substrate engagement and cleavage.

This model provides a mechanistic explanation for the switch-like activation of SAMD9 observed in cells once protein concentration exceeds a critical threshold. We further propose that pathogen-derived factors or cellular stress signals promote transient opening of SAMD9 by weakening intramolecular interactions, thereby lowering the concentration threshold required for activation. A network of individually weak, predominantly polar interactions at intramolecular interfaces, particularly those that undergo pronounced rearrangements in the asymmetric dimers, appears to support both robust autoinhibition and sensitive response to activation cues.

## Methods

### Plasmid construction

Plasmids for expression of human SAMD9 were generated by cloning full-length SAMD9 or the truncated construct encoding residues 156-1589 into the pHTN-HaloTag-CMV-neo vector (Promega, G7721). The plasmid used for transient expression of mCherry-SAMD9 fusion protein (pcDNA6.2/mCherry-SAMD9) has been described previously^28^. Site-directed mutations were introduced by assembly of PCR-amplified fragments or synthesized DNA fragments using the NEBuilder HiFi DNA Assembly Kit (NEB, E5520S). All constructs were verified by whole plasmid sequencing performed by Plasmidsaurus using Oxford Nanopore sequencing.

### Purification of Human SAMD9 Protein

Freestyle 293-F cells (ThermoFisher R79007) were maintained in vented Erlenmeyer shaker flasks (Greiner) using Freestyle 293 Expression Medium (ThermoFisher 12338-018) at 37°C with 8% CO and agitated at 120 rpm. Cells were expanded to a density of 1 × 10 cells/ml with ≥95% viability prior to transfection.

For transient expression, plasmid DNA and PEI (Polyethylenimine, linear 40 kDa; Polysciences 24765) were mixed at 1:3 (w/w) in Opti-MEM to generate DNA:PEI complexes. Complexes were added directly to Freestyle 293-F cultures at a final DNA concentration of 1 μg/ml, and protein expression proceeded for 96 h. Cells were harvested by centrifugation at 200 × g for 5 min at 4°C, washed once with DPBS, recentrifuged under the same conditions, and processed immediately for lysis.

Each ∼300-ml culture pellet was resuspended in 10 ml mammalian lysis buffer (Promega G9381) supplemented with 200 μl Protease Inhibitor Cocktail (Promega G6521), 8 μl Benzonase (Millipore 70664-3), and 50 μl of 1 M MgCl_2_. Lysates were diluted 1:3 with HaloTag Protein Purification Buffer (1 x PBS, pH 7.5, supplemented with 2 mM DTT and 0.005% IGEPAL CA-630). Clarified lysates were incubated with 500 μl HaloLink Resin (Promega G1915) pre-equilibrated in HaloTag Protein Purification Buffer. Binding was performed by inversion mixing at 4°C for ≥5 h (typically overnight). Resin was washed three times with HaloTag Protein Purification Buffer containing IGEPAL and three additional times with buffer lacking IGEPAL. SAMD9 was eluted by on-resin HaloTEV protease cleavage: 30 μl HaloTEV protease (Promega G6601) in 1.5 ml cleavage buffer was added directly to the resin, and the suspension was rotated at 4°C for ≥5 h (typically overnight). The supernatant (Elution 1) was collected by centrifugation at 1,500 × g for 5 min. Additional elutions were obtained by incubating the resin with fresh purification buffer for 30 min at 4°C.

The elutions were further purified by size-exclusion chromatography (SEC) using a Superdex 200 Increase 10/300 GL column (Cytiva) pre-equilibrated in SEC buffer (150 mM NaCl, 50 mM Tris-HCl pH 7.4, 1 mM DTT).

### Cryo-EM sample preparation, data collection, and data processing

For cryo-EM sample preparation, the SEC peak fractions of purified WT or mutant SAMD9 proteins were concentrated to ∼1 mg/mL with a 100 kDa MWCO Amicon ultrafiltration unit. An aliquot of 3.5 μL sample was applied to a glow-discharged 300-mesh Quantifoil R1.2/1.3 Cu grid, blotted with filter paper and plunge frozen in liquid ethane with Vitrobot Mark IV (ThermoFisher Scientific). Cryo-EM data collection was done with serialEM^43^ on a Titan Krios microscope equipped with Gatan BioQuantum K3 imaging filter and camera. A 20eV slit was used for the filter. Images were recorded at 130,000x magnification, corresponding to a pixel size of 0.33Å/pix at super-resolution mode of the camera. A defocus range of −1.0 to −1.8 micron was set. A total dose of 50e-/Å^2^ at dose rate of ∼13e^-^/pixel/s was fractionated into 50 frames. The first two frames of each movie stack were excluded in motion correction. A binning factor of 2 was applied during motion correction, leading to a pixel size of 0.66Å/pix for the micrographs. Cryo-EM data processing was done with cryoSPARC^44^, following standard procedures for single-particle analysis.

### Atomic modeling and refinement

The predicted structure of human SAMD9 was downloaded from the AlphaFold Protein Structure Database^45^ and fitted into the monomer density map with ChimeraX^46^ to serve as an initial model for atomic modeling. The model was refined with the Phenix real-space refinement program^47^ and manually checked in Coot^48^. This process was iterated for several cycles until no significant improvement of the model was observed. Models of the dimeric structures or mutants were modified from this monomeric model and polished in the same procedure. Structure display and figure preparation were done with ChimeraX.

### Assays of SAMD9 activity

Assays were performed as previously described^14,28^. Briefly, HEK293T cells seeded in 24-well plates were transfected with pcDNA6.2/mCherry-SAMD9 plasmids and subjected to the assays described below. To assess antiviral activity, cells transfected for 36 hours were infected with vK1^-^C7^-^/GFP^+^ VACV^49^ at an MOI of 1 for 15 hours. To measure protein synthesis, cells transfected for 24 h were incubated with 10 µM O-propargyl-puromycin (OPP) for 30 min, followed by processing with the Click-iT Plus OPP Alexa Fluor 488 Protein Synthesis Assay Kit according to the manufacturer’s instructions (ThermoFisher Scientific). To assess antiproliferative activity, cells transfected for 24 h were incubated with 10 µM 5-ethynyl-2’-deoxyuridine (EdU) for 2 h and processed using the Click-iT Plus EdU Alexa Fluor 488 Flow Cytometry Assay Kit according to manufacturer’s instructions (ThermoFisher Scientific). Total DNA was labeled with FxCycle Violet Stain (ThermoFisher Scientific). Cells were harvested using Trypsin-EDTA Solution, fixed with 4% paraformaldehyde, filtered through a 40-µm mesh, and analyzed on an LSR-II cell analyzer (BD Biosciences). Flow data were analyzed using FlowJo software (TreeStar).

### Quantification and statistical analysis

Data were analyzed using GraphPad Prism 9 software.

## Acknowledgments

The authors acknowledge the assistance of the Case Western Reserve University (CWRU) School of Medicine Cryo-Electron Microscopy Facility (supported by NIH grant S10OD032437), the CWRU High-Performance Computing Resource in the Core Facility for Advanced Research Computing, and UT Health San Antonio Flow Cytometry Shared Resource Facility (supported by NIH grants P30CA054174 and S10OD030432).

## Author contributions

Conceptualization: Y.X. and X.D. Methodology: Z.M, F.Z. Investigation: Z.M, F.Z., M.M., B.S. Visualization: Z.M, F.Z., M.M., B.S., X.D., and Y.X. Funding acquisition: Y.X., and X.D. Project administration: Y.X., and X.D. Supervision: Y.X., and X.D. Writing—original draft: Y.X. and X.D. Writing—review and editing: Z.M, F.Z., X.D., and Y.X.

## Data, code, and materials availability

All data are available in the main text or the supplementary materials. All atomic models and cryo-EM maps have been deposited in the Protein Data Bank (PDB) and Electron Microscopy Data Bank (EMDB), respectively, with the following identifiers: monomer, 9ZJR, EMD-74339; symmetric dimer, 9ZJU, EMD-74342; asymmetric dimer “shell” shape, 9ZJS, EMD-74340; asymmetric dimer “wing” shape, 9ZJV, EMD-74343; LoF mutant R837A, 9ZJW, EMD-74344; LoF mutant R685Q&G686C&I968C, 9ZJZ, EMD-74347. The materials used in the paper can be obtained from the corresponding authors pending a completed material transfer agreement.

