## Supplementary material for "Structural Mechanisms of SAMD9 Autoinhibition and Pathogenic Dysregulation": Figs. S1 to S10,Tables S1 to S2, and Supplemental file 1

**Supplementary Materials**

Figs. S1 to S10

Tables S1 to S2

Supplemental file 1

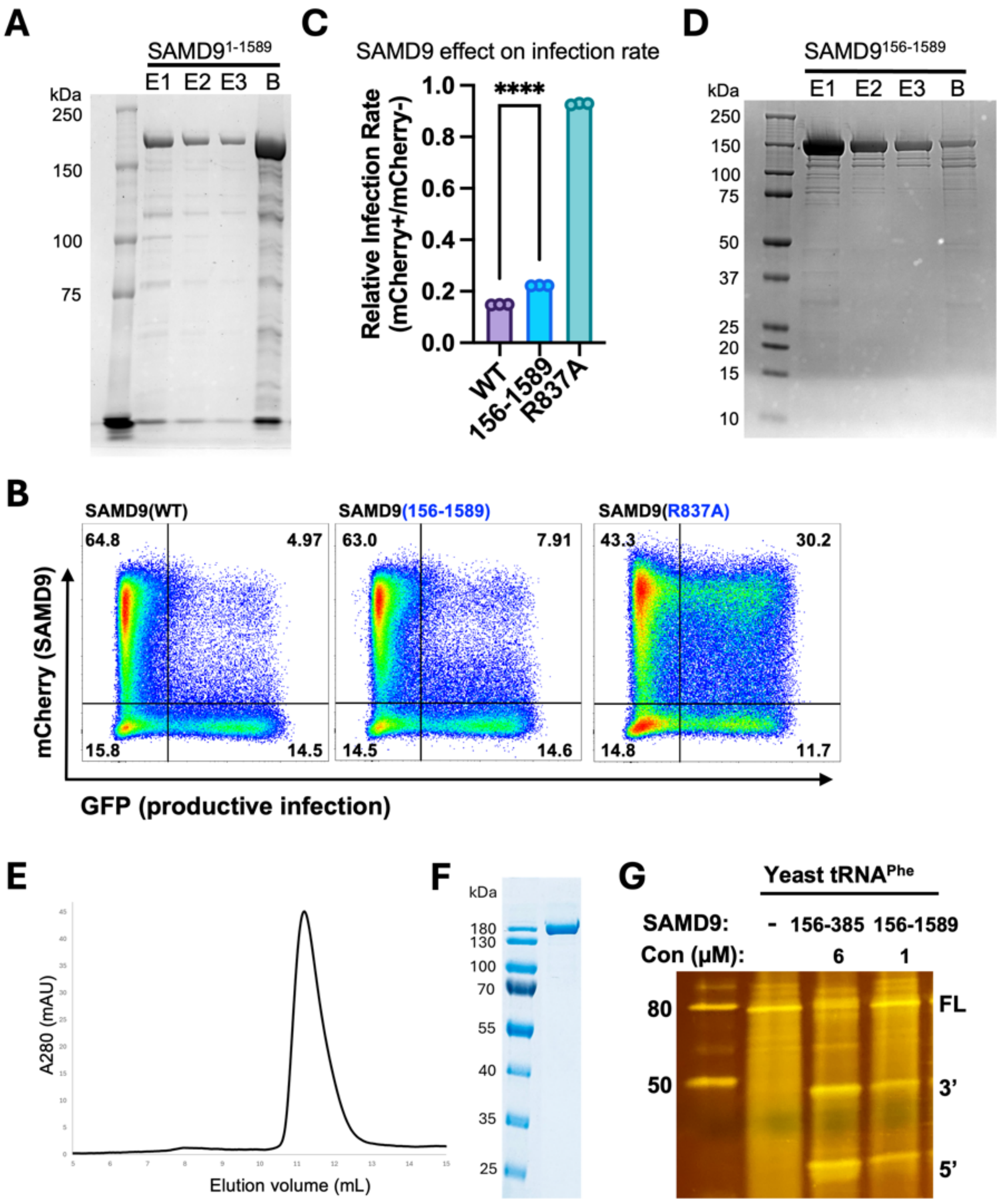


**Figure S1. Characterization and purification of SAMD9.**

**(A)** Coomassie-stained SDS-PAGE gel shows the purification of full-length SAMD9 protein. SAMD9 with an N-terminal HaloTag followed by a TEV protease cleavage site was transiently expressed in HEK293-F cells. Clarified cell lysates were incubated with HaloLink resin, and bound SAMD9 was released by on-resin HaloTEV protease cleavage. Shown are three elution fractions following HaloTag cleavage (E1, E2, and E3), along with residual protein remaining on the HaloLink resin beads (B). **(B)** HEK293T cells were transfected with mCherry-SAMD9 fusion constructs and infected with vK1$^{-}$C7$^{-}$/GFP$^{+}$ VACV. Representative flow-cytometry plots show virus-encoded GFP expression versus mCherry expression, with quadrant percentages indicated. **(C)** Quantification of relative VACV infection rates in SAMD9-expressing versus non-transfected cells within the same sample. **(D-F)** Expression and purification of SAMD9$^{156\text{-}1589}$ from HEK293-F cells. **(D)** Coomassie-stained SDS-PAGE gel showing three elution fractions following HaloTag cleavage and residuals on the beads. **(E)** Size-exclusion chromatography (SEC) profile of Elution 1 from panel D. **(F)** Coomassie-stained SDS-PAGE gel of the final purified SAMD9$^{156\text{-}1589}$ protein following SEC. **(G)** In vitro tRNase assay. Purified yeast tRNA$^{Phe}$ was incubated with the indicated proteins for 1 h, and cleavage products were visualized by SYBR Gold staining. FL denotes full-length tRNA$^{Phe}$; 3′ and 5′ indicate the corresponding cleavage products. "–" indicates no protein added.

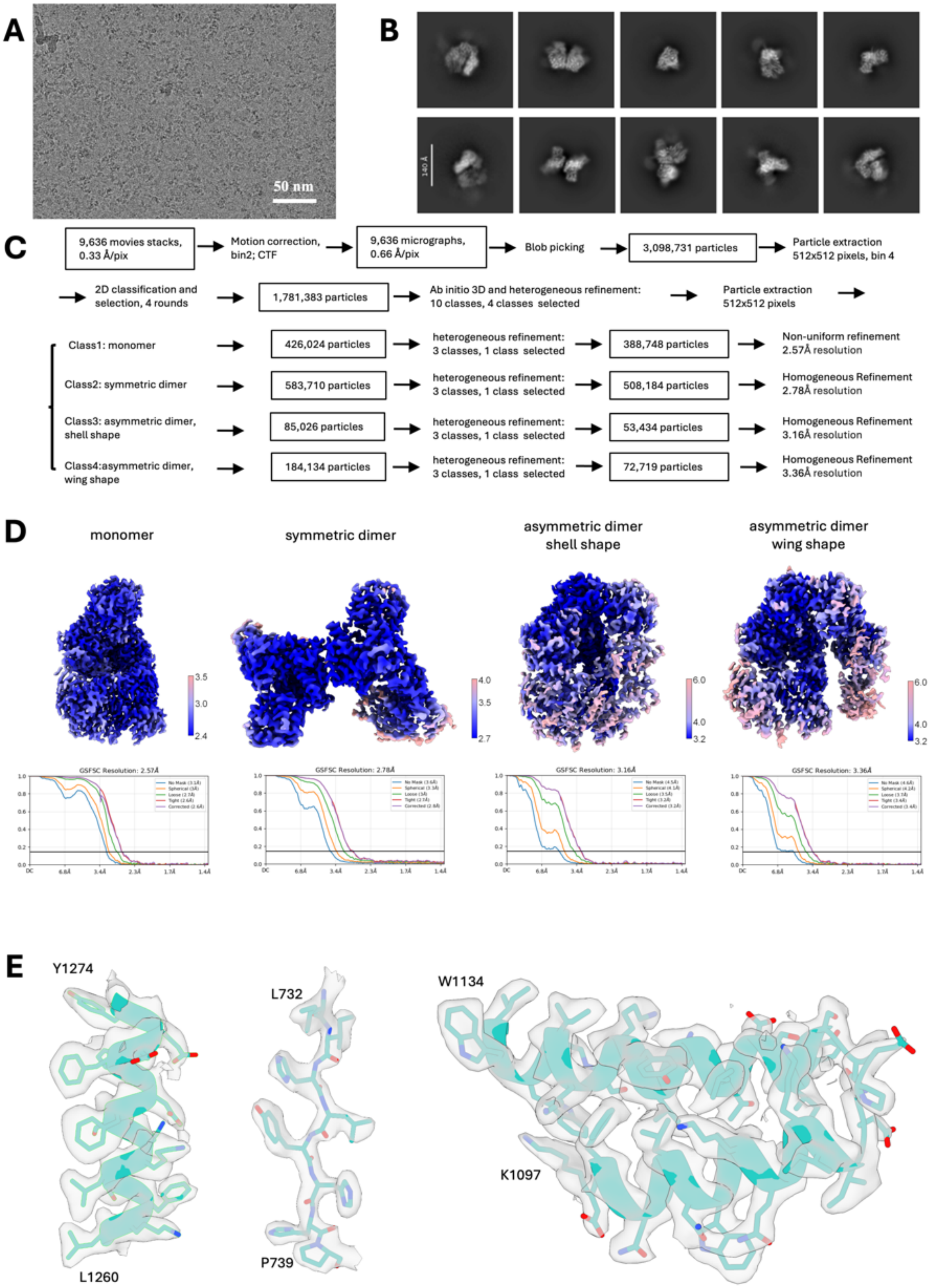


**Figure S2. Cryo-EM data collection, processing, and quality assessment of SAMD9 structures.**
**(A)** Representative cryo-EM micrograph of purified human SAMD9$^{156\text{-}1589}$ particles. Scale bar, 50 nm. **(B)** Representative two-dimensional (2D) class averages of SAMD9 particles. **(C)** Cryo-EM data-processing workflow

leading to the determination of SAMD9 structures. **(D)** Local resolution maps and corresponding gold-standard Fourier shell correlation (GSFSC) curves for the SAMD9 monomer, C2-symmetric dimer, asymmetric shell dimer, and asymmetric wing dimer, shown from left to right. **(E)** Segmented cryo-EM density maps with their corresponding atomic models, highlighting high-resolution features of the reconstructed structures. For clarity, only the N- and C-terminal residues of each modeled segment are labeled.

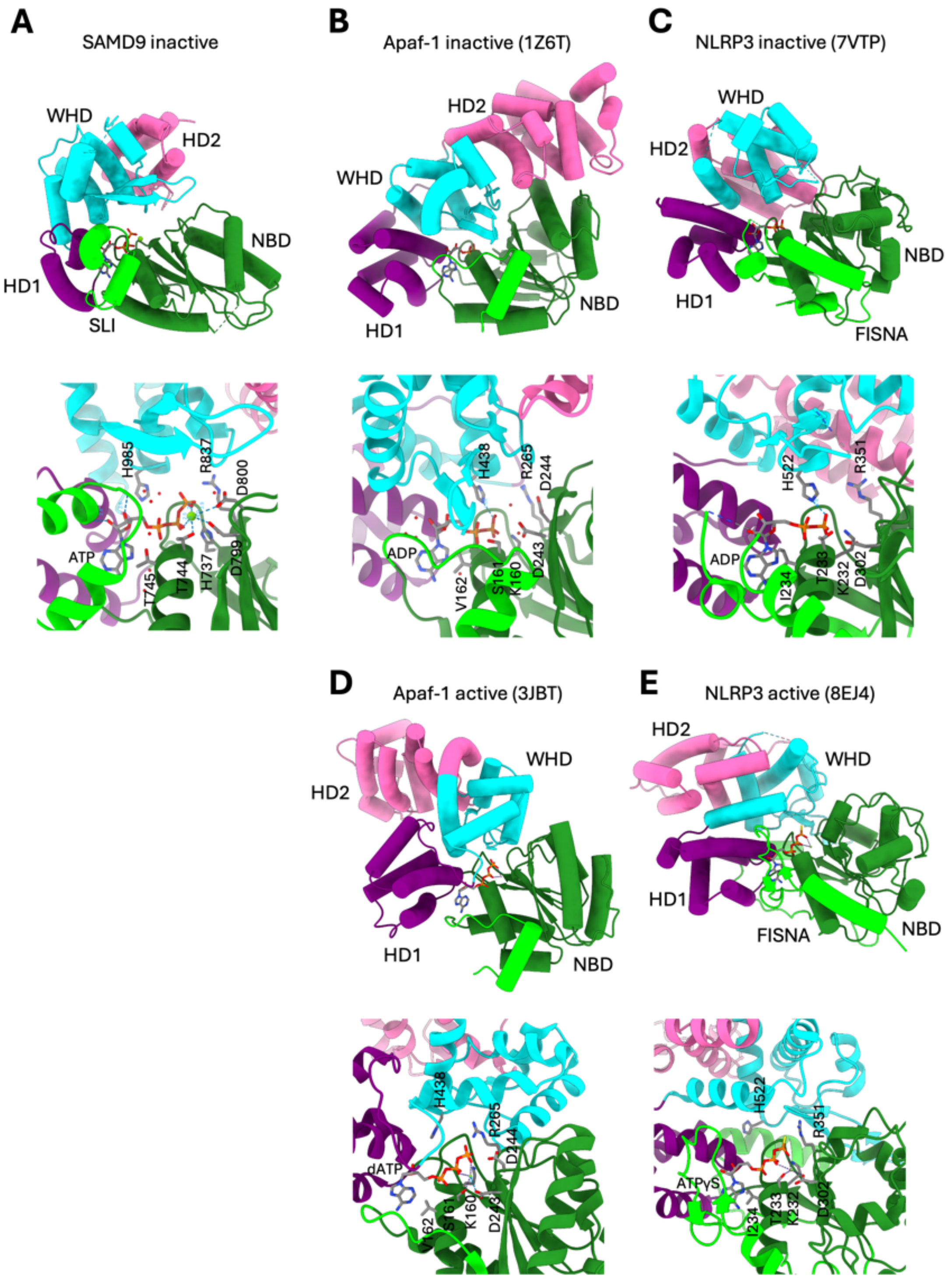


**Figure S3. Structural comparison of the SAMD9 NOD module with those of Apaf-1 and NLRP3.** **(A-C)** The NOD modules of SAMD9 (A), Apaf-1 (B), and NLRP3 (C) in their autoinhibited state are depicted side by side in tube-helix mode. Bound nucleotides (ATP for SAMD9; ADP for Apaf-1 and NLRP3) are displayed as sticks. Conserved NOD domains, including the NBD, HD1, WHD, and HD2, are labeled. The lineage-specific N-terminal extensions immediately preceding the NBD are highlighted in lime: the SAMD9-Lineage Insert (SLI) in SAMD9 and the FISNA motif in NLRP3. Notably, SAMD9 displays a more pronounced cleft within the NOD module compared with Apaf-1 and NLRP3. The conserved WHD histidine contacts the β-phosphate of ADP in Apaf-1 and NLRP3 but is positioned slightly further from the ATP phosphates in SAMD9. **(D-E)** NOD modules of

Apaf-1 (D) and NLRP3 (E) in their ATP-bound, active states, illustrating the large rigid-body rotation of the WHD-HD2 submodule away from the nucleotide-binding pocket.

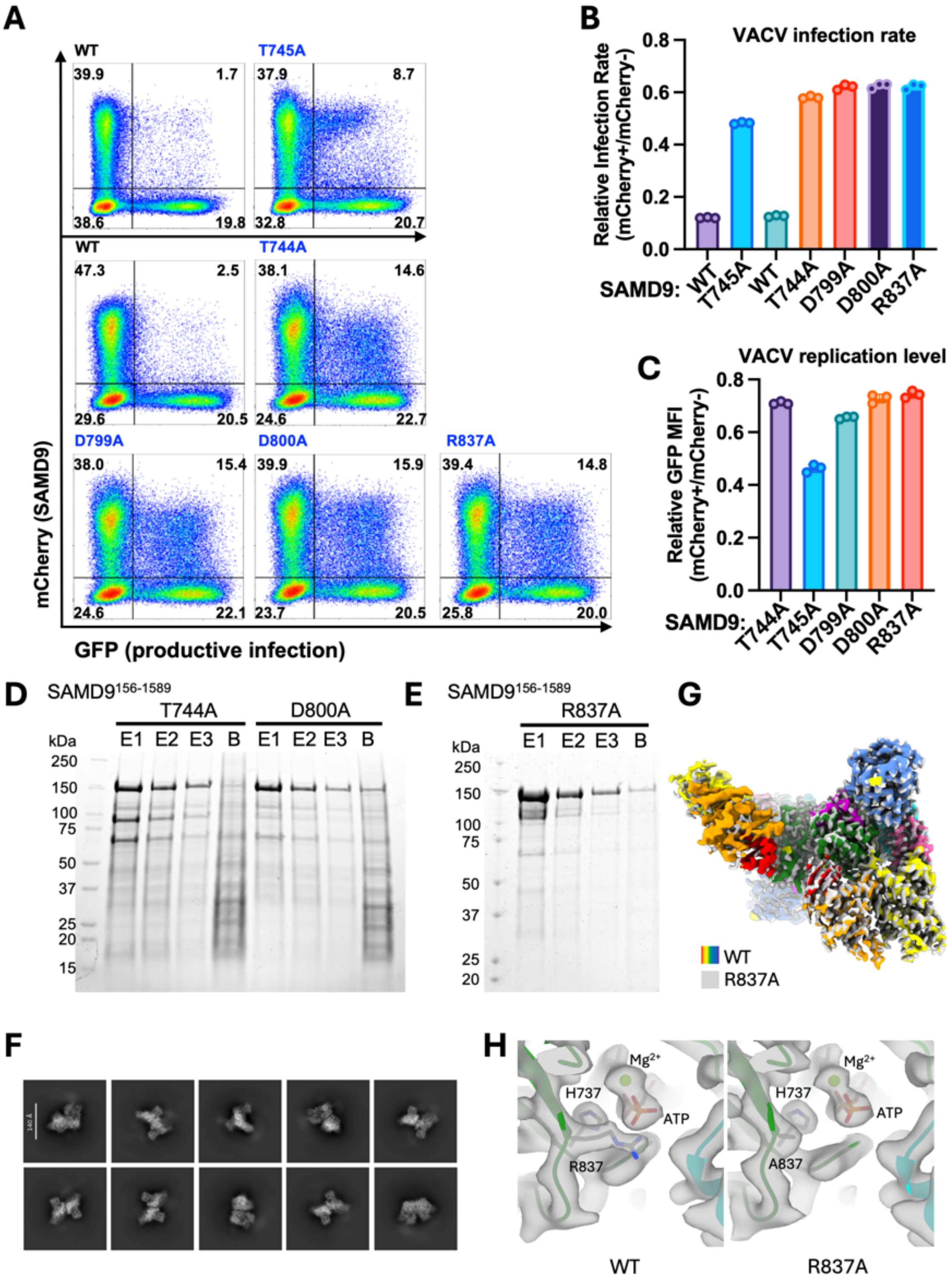


**Figure S4. Mutations of ATP-contacting residues cause loss of function.**
(**A**) HEK293T cells were transfected with mCherry-SAMD9 fusion constructs and infected with vK1$^{-}$C7$^{-}$/GFP$^{+}$ VACV. Representative flow-cytometry plots show virus-encoded GFP expression versus mCherry expression, with quadrant percentages indicated. **(B)** Quantification of relative VACV infection rates (% of GFP$^{+}$ cells) in SAMD9-expressing versus non-transfected cells within the same sample. **(C)** Quantification of relative VACV replication level (GFP MFI) in SAMD9-expressing versus non-transfected cells within the same sample. Note that T745A exhibits a milder defect compared with the other mutants, as indicated by a more pronounced reduction in viral

infection rate and replication level. **(D-E)** Expression and purification of T744A, D800A (D) and R837A (E) variants of SAMD9$^{156\text{-}1589}$ from HEK293-F cells. Shown are three elution fractions following HaloTag cleavage, along with residual protein remaining on the HaloLink resin beads. **(F)** Representative cryo-EM 2D class averages of SAMD9 R837A mutant particles. The sample is dominated by C2 symmetric dimers compared to the WT in Fig. S2B. (**G**) Cryo-EM density map of SAMD9 R837A (grey) superimposed with that of the WT (domain colored as in Fig. 1C). The two structures are nearly identical. (**H**) Side-by-side comparison of the sidechain densities at the R837A mutation site (right) with that of the WT (left). Note that the mutation did not affect ATP binding.

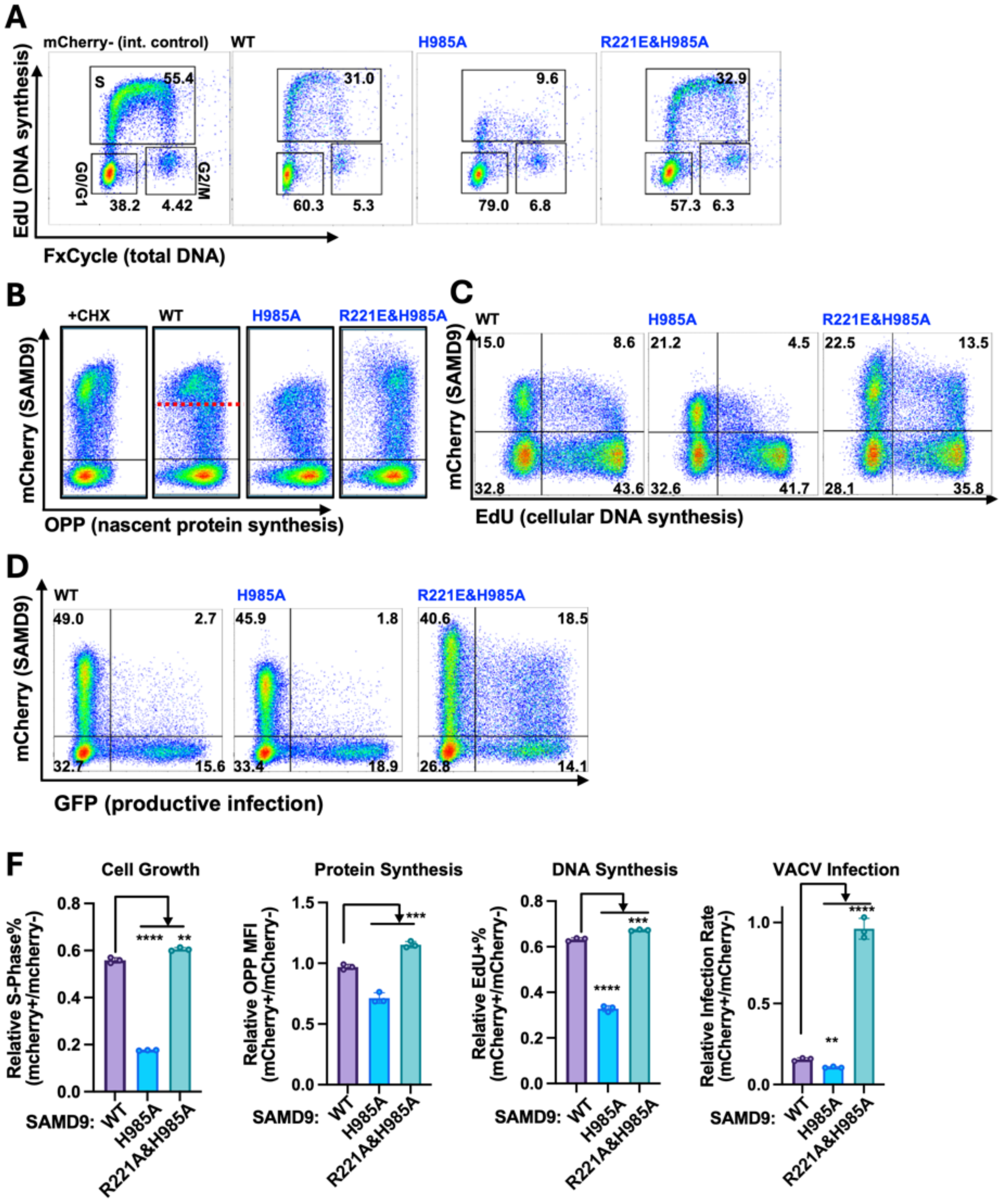


**Figure S5. Mutation of WHD H985 causes spontaneous activation.**
**(A-D)** HEK293T cells were transfected with mCherry-SAMD9 fusion constructs. **(A)** Cells were labeled with FxCycle and EdU (2 h pulse) to measure total cellular DNA and newly synthesized DNA, respectively. Representative flow cytometry plots show cellular EdU and FxCycle levels amongst mCherry-SAMD9$^-$ or mCherry-SAMD9$^+$ population. **(B)** Cells were labeled with O-propargyl-puromycin (OPP) for 30 min to measure nascent protein synthesis. Representative flow-cytometry plots show OPP incorporation versus mCherry expression. Untransfected cells in the presence of cycloheximide (CHX) are shown as a control. The red dashed line indicates the mCherry-SAMD9 expression threshold above which inhibition of protein synthesis is observed. **(C)** Cells were labeled with EdU (2 h pulse) to assess DNA synthesis. Representative flow-cytometry plots show EdU incorporation versus mCherry expression. **(D)** Cells were infected with a SAMD9-susceptible VACV mutant, vK1$^-$C7$^-$/GFP$^+$. Representative flow-cytometry plots show virus-encoded GFP expression versus mCherry expression, with quadrant

percentages indicated. **(E)** Relative S-phase fractions, nascent protein synthesis, cellular DNA synthesis, and infection rates in SAMD9-expressing versus non-transfected cells within the same sample were quantified from the flow-cytometry data in panels **A-D**. Statistical significance was assessed by pairwise Student's *t* tests (****, $P < 0.0001$; ***, $P < 0.001$;**, $P < 0.01$).

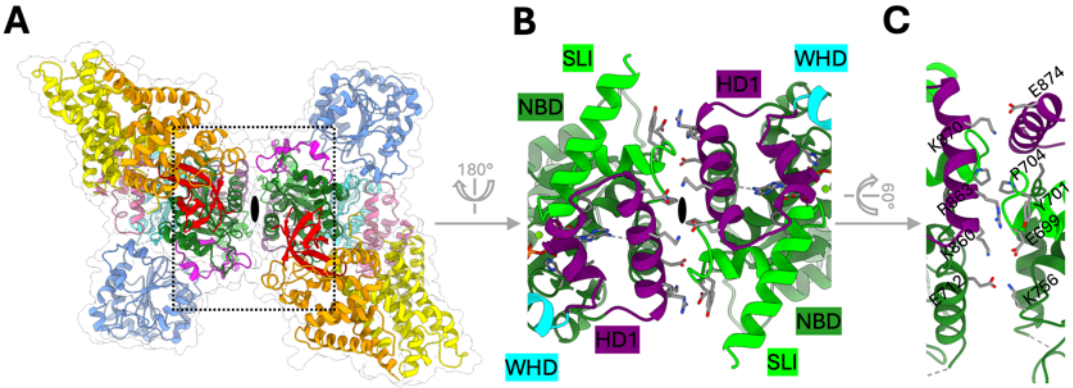


**Figure S6. Intermolecular interactions in the C2-symmetric SAMD9 dimer.**
**(A)** Atomic model of the SAMD9 C2-symmetric dimer shown within a semi-transparent white envelope representing the overall particle shape. The black oval denotes the C2 symmetry axis, which is perpendicular to the plane of the dimer interface. Individual domains in each protomer are color-coded as in Figure 1B. **(B, C)** Close-up views of the intermolecular dimerization interface. The region shown in (B) corresponds approximately to the dotted box in (A), viewed from the opposite side of the dimer. In (C), the structure is rotated to highlight key sidechain interactions across the interface; for clarity, the symmetry-related set of sidechains from the opposing protomer is omitted.

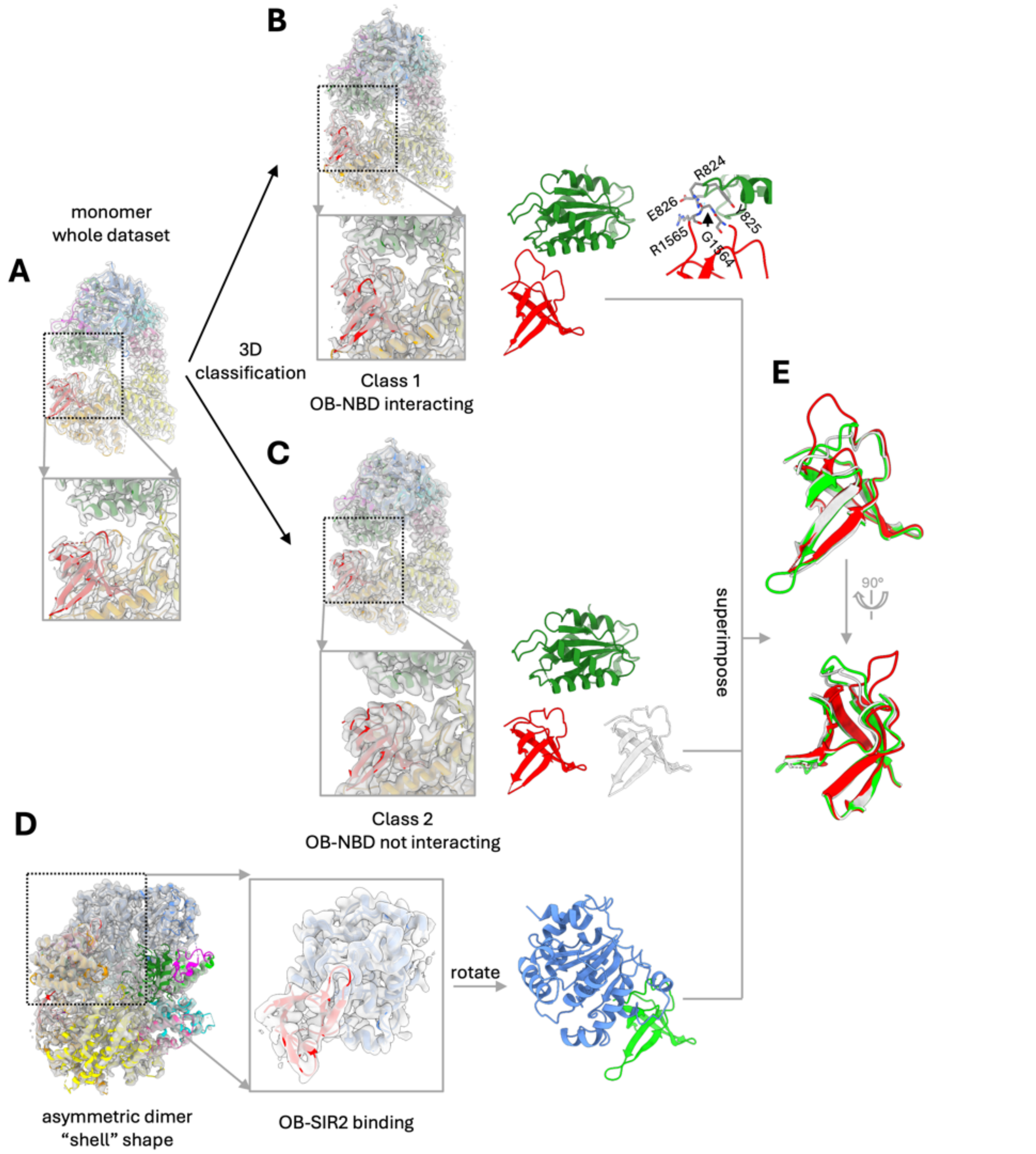


**Figure. S7. The structural dynamics of OB/NBD and OB/SIR2 interactions.**
**(A-C)** 3D classification focusing on the TPR2-OB and NBD regions in the SAMD9 monomer particles reveals the dynamic nature of OB/NBD interactions. In the reconstruction of SAMD9 monomer with the whole dataset, OB appears separated from the NBD (A). After focused 3D classification, two major classes emerged. In class 1, an OB loop is extended out to engage NBD (B), and in class 2, the OB loop stays away from the NBD (C). **(D)** Structure of the OB/SIR2 in the asymmetric shell dimer. **(E)** OB models in the NBD-engaged (red), NBD-disengaged (white) or SIR2-engaged (green) states are superposed and compared. The similarity between NBD-disengaged and SIR2-engaged OB structures suggests that disengaging from the NBD primes OB for SIR2 binding.

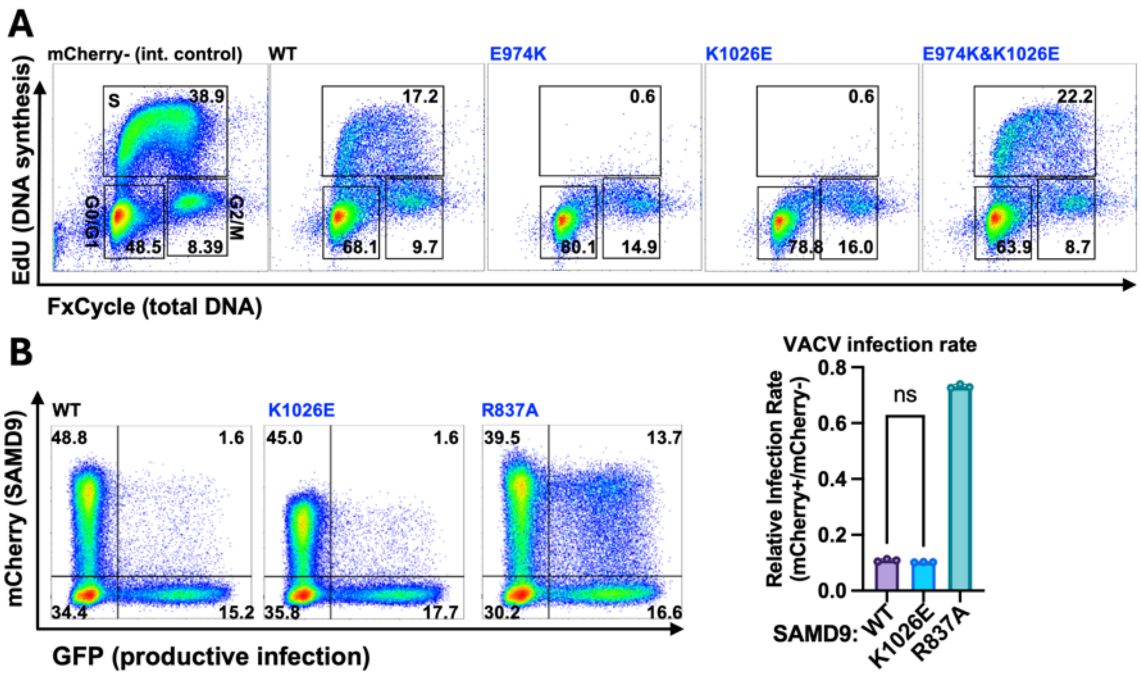


**Figure S8. Restoration of the SIR2/(WHD-HD2) interface suppresses pathological activation by the GoF E974K mutation.**

**(A)** HEK293T cells were transfected with mCherry-SAMD9 fusion constructs and labeled with FxCycle and EdU (2 h pulse) to measure total cellular DNA and newly synthesized DNA, respectively. Representative flow cytometry plots of cellular EdU and FxCycle levels amongst mCherry-SAMD9$^{+}$ or mCherry-SAMD9$^{-}$ population are shown. The derived S-phase levels in SAMD9-expressing cells relative to non-transfected cells from the same culture wells are shown in Fig. 3E. **(B)** HEK293T cells were transfected with mCherry-SAMD9 fusion constructs and infected with vK1$^{-}$C7$^{-}$/GFP$^{+}$ VACV. Representative flow-cytometry plots show virus-encoded GFP expression versus mCherry expression, with quadrant percentages indicated. Relative infection rates in SAMD9-expressing versus non-transfected cells from the same culture well were quantified. Statistical significance was determined using pairwise Student's *t* tests (ns, not significant).

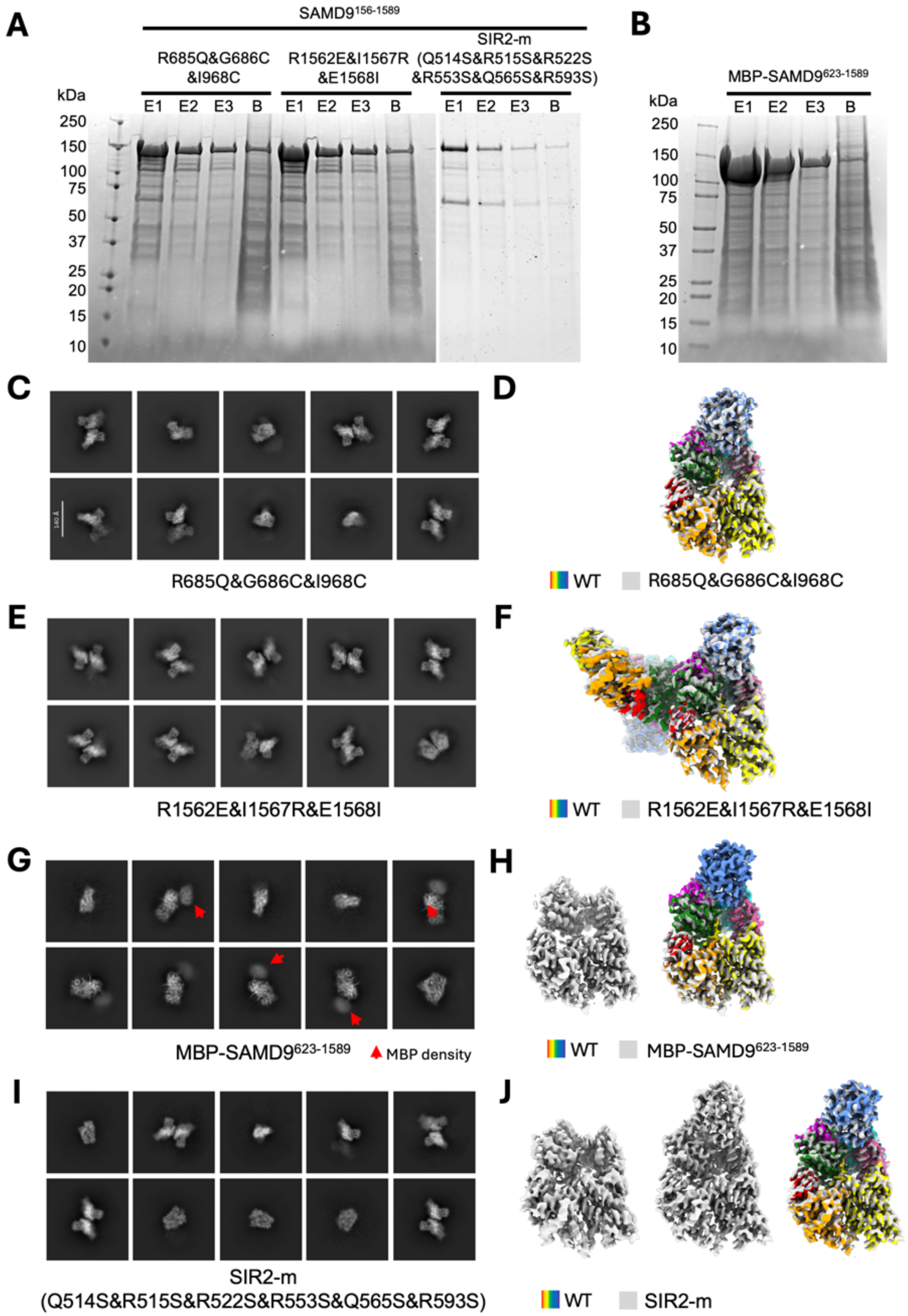


**Figure S9. Cryo-EM structures of SAMD9 mutants.**

(**A**, **B**) Expression and purification of SAMD9 proteins from HEK293-F cells. Shown are three elution fractions following HaloTag cleavage, along with residual protein remaining on the HaloLink resin beads. (**C**, **E, G, I**) Representative cryo-EM 2D class averages of SAMD9 mutants as indicated below each panel. Red arrows in (G) point to fuzzy MBP densities. (**D**, **F, H, J)** Cryo-EM density maps of SAMD9 mutants (grey) superimposed with that of the WT (domain colored as in Fig. 1C) to show that the mutations did not change the overall structure of the closed state. The MBP moiety in MBP-SAMD9$^{623\text{-}1589}$ construct is flexible and not resolved as shown in (H). The SIR2 domain in a subset of Sir2-m particles is also flexible and not resolved as shown in (J).

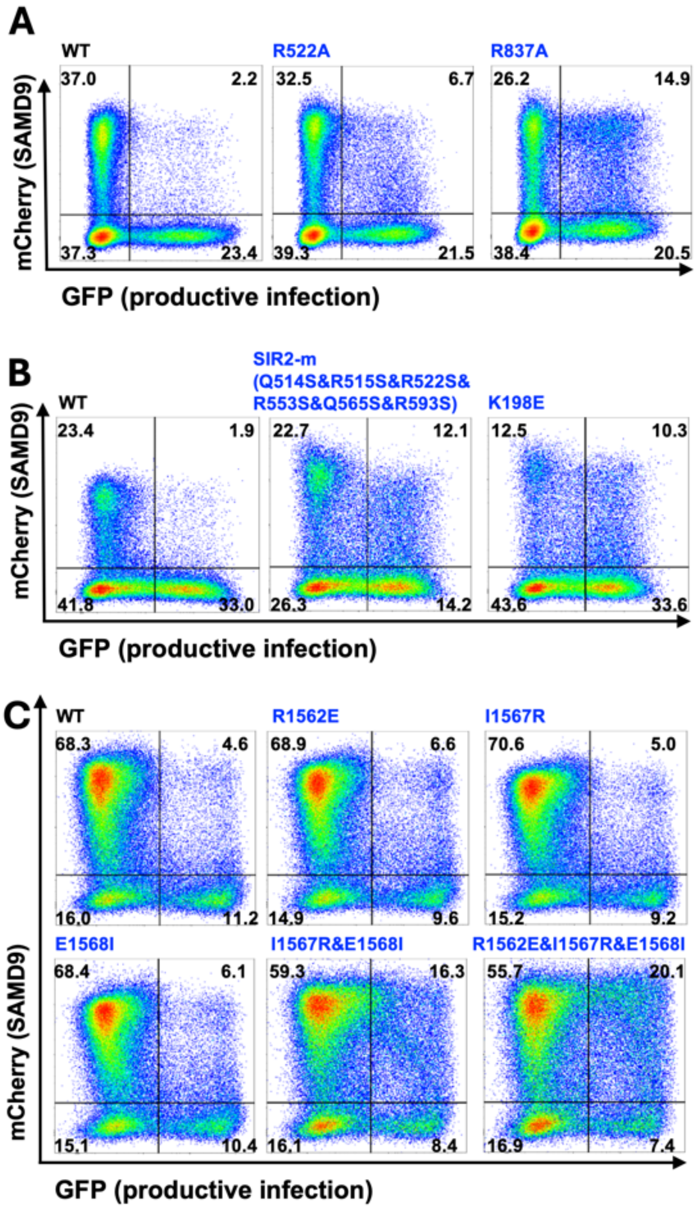

**Figure S10. Intermolecular SIR2/SIR2 and OB/SIR2 contacts are critical for SAMD9 activation.** HEK293T cells were transfected with mCherry-SAMD9 fusion constructs and infected with vK1$^{-}$C7$^{-}$/GFP$^{+}$ VACV. Representative flow-cytometry plots show virus-encoded GFP expression versus mCherry expression, with quadrant percentages indicated. Quantifications of relative infection rates in SAMD9-expressing cells versus mCherry-negative cells are shown in Fig. 5H and Fig. 5J. **(A, B)** Mutations targeting the SIR2/SIR2 dimerization interface. **(C)** Mutations targeting the OB/SIR2 interaction interface.

### Table S1. Cryo-EM data collection, refinement and validation statistics

| | #1 monomer (EMDB 74339) (PDB 9ZJR) | #2 C2 symmetric dimer (EMDB 74342) (PDB 9ZJU) | #3 Asymmetric shell dimer (EMDB 74340) (PDB 9ZJS) |
|---|---|---|---|
| **Data collection and processing** | | | |
| Magnification | 130,000x | 130,000x | 130,000x |
| Voltage (kV) | 300 | 300 | 300 |
| Electron exposure (e–/Å$^2$) | 50 | 50 | 50 |
| Defocus range (µm) | -0.8 to -1.5 | -0.8 to -1.5 | -0.8 to -1.5 |
| Pixel size (Å) | 0.66 | 0.66 | 0.66 |
| Symmetry imposed | C1 | C1 | C1 |
| Initial particle images (no.) | 3,098,731 | 3,098,731 | 3,098,731 |
| Final particle images (no.) | 388,748 | 508,184 | 54,343 |
| Map resolution (Å) | 2.57 | 2.78 | 3.16 |
| FSC threshold | 0.143 | 0.143 | 0.143 |
| Map resolution range (Å) | 2.5-4.0 | 2.7-4.0 | 3-4.0 |
| **Refinement** | | | |
| Initial model used (PDB code) | | | |
| Model resolution (Å) | 2.57 | 2.78 | 3.16 |
| FSC threshold | 0.143 | 0.143 | 0.143 |
| Model resolution range (Å) | 2.5-4.0 | 2.7-4.0 | 3-4.0 |
| Map sharpening *B* factor (Å$^2$) | 92.6 | 86.9 | 64.2 |
| Model composition | | | |
| Non-hydrogen atoms | 9391 | 18426 | 17812 |
| Protein residues | 1121 | 2242 | 2164 |
| Ligands | 2 | 4 | 4 |
| *B* factors (Å$^2$) | | | |
| Protein | 6.5/117.6/46.4 | 22.6/193/90.9 | 15.06/217.98/108.67 |
| Ligand | 9.93/35.6/17.5 | 25.9/54.4/37.8 | 40.33/153.81/9783 |
| R.m.s. deviations | | | |
| Bond lengths (Å) | 0.003 | 0.002 | 0.004 |
| Bond angles (°) | 0.453 | 0.471 | 0.525 |
| Validation | | | |
| MolProbity score | 1.7 | 1.43 | 2.38 |
| Clashscore | 4.01 | 4.66 | 25.91 |
| Poor rotamers (%) | 3.13 | 1.76 | 2.63 |
| Ramachandran plot | | | |
| Favored (%) | 97.27 | 98.36 | 97.15 |
| Allowed (%) | 2.73 | 1.64 | 2.85 |
| Disallowed (%) | 0.00 | 0.00 | 0.00 |

### Cryo-EM data collection, refinement and validation statistics

| | #4 Asymmetric wing dimer (EMDB 74343) (PDB 9ZJV) | #5 Mutant R685Q&G686C&I968C (EMDB 74347) (PDB 9ZJZ) | #6 Mutant R837A (EMDB 74344) (PDB 9ZJW) |
|---|---|---|---|
| **Data collection and processing** | | | |
| Magnification | 130,000x | 130,000x | 130,000x |
| Voltage (kV) | 300 | 300 | 300 |
| Electron exposure (e–/Å$^2$) | 50 | 50 | 50 |
| Defocus range (μm) | -0.8 to -1.5 | -0.8 to -1.5 | -0.8 to -1.5 |
| Pixel size (Å) | 0.66 | 0.66 | 0.66 |
| Symmetry imposed | C1 | C1 | C1 |
| Initial particle images (no.) | 3,098,731 | 1,279,061 | 2,119,769 |
| Final particle images (no.) | 72,719 | 165,609 | 383,351 |
| Map resolution (Å) | 3.36 | 2.87 | 2.70 |
| FSC threshold | 0.143 | 0.143 | 0.143 |
| Map resolution range (Å) | 3.3-4.0 | 2.9-4.0 | 2.6-4.0 |
| **Refinement** | | | |
| Initial model used (PDB code) | | | |
| Model resolution (Å) | 3.36 | 2.87 | 2.70 |
| FSC threshold | 0.143 | 0.143 | 0.143 |
| Model resolution range (Å) | 3.3-4.0 | 2.9-4.0 | 2.6-4.0 |
| Map sharpening *B* factor (Å$^2$) | 67.6 | 90.6 | 86.2 |
| Model composition | | | |
| Non-hydrogen atoms | 14797 | 9213 | 18426 |
| Protein residues | 1804 | 1121 | 2242 |
| Ligands | 2 | 2 | 4 |
| *B* factors (Å$^2$) | | | |
| Protein | 33.45/213.4/112.92 | 32.73/163.91/74.8 | 68.9/385.02/162.97 |
| Ligand | 59.51/96.43/71.04 | 39.94/80.23/54.95 | 79.11/121.36/94.07 |
| R.m.s. deviations | | | |
| Bond lengths (Å) | 0.003 | 0.003 | 0.003 |
| Bond angles (°) | 0.569 | 0.514 | 0.582 |
| Validation | | | |
| MolProbity score | 1.72 | 1.61 | 1.51 |
| Clashscore | 7.52 | 5.80 | 7.13 |
| Poor rotamers (%) | 1.89 | 2.45 | 1.42 |
| Ramachandran plot | | | |
| Favored (%) | 97.51 | 98.27 | 98.04 |
| Allowed (%) | 2.49 | 1.73 | 1.92 |
| Disallowed (%) | 0.00 | 0.00 | 0.00 |

**Table S2. Gain-of-function (GoF) SAMD9 mutations identified in patients and their possible structural explanations.**

| Mutation | Domain | Affected domain or domain-domain interface | Possible structural explanation |
|---|---|---|---|
| F437S | SIR2 | SIR2/NOD | F437 is stacking with L485 & M534, both of which are located on a SIR2 helix or loop that are interacting with NOD; therefore, F437S likely destabilizes the SIR2/NOD interface |
| R459Q[a] | SIR2 | SIR2/Δ | directly located at the SIR2/Δ interface; the sidechain of R459 has a fixed conformation by forming hydrogen bond with the carbonyl of K432, as well as stacking interactions with K432, Y455 (both in SIR2), and K643 (on Δ). R459Q likely disrupts this intricate network of interactions, weakening the SIR2/Δ-NOD interface. |
| F465L | SIR2 | SIR2/NOD | F465 is stacking with Y505, and has hydrophobic interactions with several other neighboring residues, "hooking" the segment 461-489 in position. This segment contains the short helix 476-489 that is involved in SIR2/NOD interaction. |
| Y469C/S | SIR2 | SIR2/NOD | Y469 has hydrophobic/stacking interaction with L483 to stabilize the helix 476-489 that is involved in SIR2/NOD interaction. |
| L483V | SIR2 | SIR2/NOD | L483 has stacking interaction with Y469 and H464 to stabilize the helix 476-489 that is involved in SIR2/NOD interaction. |
| N484S | SIR2 | SIR2/Δ | N484 is at the interface of SIR2/Δ. |
| Y486S | SIR2 | SIR2/NOD | Y486 stacking with the mainchain of G976 seems to be a major interaction between SIR2 and the beta-turn-beta structure of WHD. |
| L641P | Δ | SIR2/Δ | L641P might affect the folding of the Δ domain, and thus the SIR2/Δ interaction |
| K643E | Δ | SIR2/Δ | As explained for R459Q; K643/R459 interaction is at the SIR2/Δ interface. |
| N658D | Δ | SIR2/Δ | The sidechain of N658 forms two hydrogen bonds with the backbone of 629 and 631, stabilizing this beta-strand of the Δ domain. N658D might also have repelling charge-charge interaction with neighboring E657 and E661, further destabilizing the Δ fold. |
| D668N | SLI | Δ/SLI | D668 is at the boundary of two helices connecting Δ to SLI, where a nearly 90-degree turn is observed between the two helices. D668N might have affected the folding of the two helices. |

| | | | |
|---|---|---|---|
| K676E | SLI | SLI | K676 has charge-charge interaction with E680 on the same helix and stacking interaction with F696 on the other helix of SLI. K676E may change the conformation of the two helices of SLI. |
| R685Q | SLI | SLI/WHD | As explained in the main text, R685 is forming hydrogen bonds with the mainchain of 964 & 967 in the WHD, stabilizing the NOD module. |
| V689A/L | SLI | SLI, ATP binding? | V689 sidechain is inserted into a hydrophobic pocket formed by F663, F705, F694, M748; its mainchain amide also forms a hydrogen bond with a water molecule, which in turn interacts with the α-phosphate of the bound ATP. V689A/L with a shorter or longer sidechain may change the intricate balance of these interactions. |
| R708G | NBD | SLI, NBD, HD1 | R708 is at the boundary of SLI, NBD and HD1 domains; its sidechain forms hydrogen bonds with the carbonyl of I854, C741, and a water molecule. R708G might have affected the relative positioning of SLI, NBD, and HD1. |
| E712K | NBD | C2 dimerization? | Not clear. E712/K756 is one of the charge-charge interactions leading to C2 dimerization, which stabilizes the autoinhibited state as explained in the main text. E712K may prevent C2 dimerization. |
| L714P | NBD | NBD | L714P is in an α-helix of NBD, L714P likely will end the α--helix folding at this position, making the helix shorter. |
| A722E | NBD | NBD | A722 is facing hydrophobic residues I831, F758, P793 of NBD. A722E may change the conformation of the helix 709-723 in NBD. |
| I750F | NBD | NBD | Similar to A722E, the bulkier I750F may push the neighboring helix 709-723 away, affecting the conformation of the NBD. |
| T767K/I | NBD | SIR2/NOD | T767 is at the boundary of SIR2, NBD and WHD. |
| D769G/N | NBD | NBD/TPR1 | Might affect the binding of the loop extended from TPR1, which stabilizes the autoinhibited state. |
| I773M/T | NBD | NBD | I773 is a buried hydrophobic residue. I773M/T might affect the conformation of the α-helix 769-782 in NBD. |
| Q776K | NBD | SIR2/Δ | Q776 has stacking and hydrogen bond interactions with M649, T650, A651 of Δ. |
| T778I | NBD | NBD | T778I may affect NBD conformation locally. |
| E803Q | NBD | NBD/TPR1 | E803 has charge-charge interaction with R1188 on the loop extended from TRP1 to NBD, which stabilizes the autoinhibited state. |
| V807I | NBD | NBD | V807 is buried with the NBD. V807I may affect NBD conformation locally. |
| K821M | NBD | NBD | K821 is interacting with E775 within NBD. K821M may affect NBD conformation locally. |
| R824Q[a] | NBD | NBD/OB | R824 is likely involved in binding the OB loop 1556-1568, stabilizing the autoinhibited state. |

| | | | |
|---|---|---|---|
| N834Y | NBD | NBD | N834 is buried within NBD. N834Y may affect NBD conformation locally. |
| H876R | HD1 | HD1/SLI | The sidechain of H876 is stacking with that of F882, maintaining the highly skewed $3_{10}$ helix 876-882 that connects the other two regular α-helices in HD1. H876R may affect the conformation of HD1 and its interaction with SLI. |
| K877E | HD1 | HD1 | similar to H876R, K877E may affect the conformation of the $3_{10}$ helix 876-882 of HD1. |
| I897M | WHD | HD1/WHD | I897 is buried and involved in hydrophobic interaction between HD1 and WHD. |
| R902W | WHD | WHD | R902 interacts locally with E898 and E995 within WHD. |
| L924V | WHD | WHD/HD2 | L924 is buried and involved in hydrophobic interaction with HD2. |
| G952R | WHD | SIR2/NOD | G952 is on a relatively flexible loop 945-954 that extends from the WHD to interact with SIR2. W951 on this loop likely stacks with Y409 on SIR2, stabilizing the SIR2/NOD interaction. G952R may render this loop less flexible, affecting its binding with SIR2. |
| V972F | WHD | SIR2/NOD | V972 is on the beta-turn-beta structure of WHD that is involved in interaction with SIR2. |
| E974K[a] | WHD | HD2 | Similar to V972F, E974K may affect the conformation of the beta-turn-beta structure of the WHD that is involved in interaction with SIR2. E974K likely will have repelling charge-charge interaction with K1026. |
| R982H/C[a] | WHD | WHD/SLI | R982 forms hydrogen bonds with the backbone of SLI and water molecules near the bound ATP. R982H/C likely will affect the stability of the NOD. |
| I983V/S | WHD | NOD | I983 sidechain is embedded within WHD, but its mainchain carbonyl is involved in hydrogen bonds with ordered water molecules bridging the two sides of the NOD module. I983V/S might affect the stability of the NOD module. |
| I988M | WHD | HD1/WHD | Similar to I897M. I988 is next to I897 and I887, involved in hydrophobic interactions between HD1 and WHD. |
| N1003K | HD2 | HD2/TPR1? | Not clear. N1003K might attract neighboring E1062 and D1061, affecting the conformation of HD2 and its interface with TPR1. |
| E1014K | HD2 | WHD/HD2 | E1014K might repel neighboring K913 and K916 in WHD and destabilize the WHD/HD2 interface. |
| F1017V | HD2 | WHD/HD2 | F1017 interacts with several other hydrophobic residues at the WHD/HD2 interface. F1017V may destabilize the WHD/HD2 interface with weakened hydrophobic interaction. |
| S1074I | HD2 | HD2/TPR1 | S1074 is surrounded by several hydrophobic residues within HD2. S1074I may change the conformation of helix 1064-1078 and affect the HD2/TPR1 interface. |

| | | | |
|---|---|---|---|
| F1092L | TPR1 | HD2/TPR1 | F1092 is mediating a series of stacking interactions among N1101, F1092, H1091, H1059, V1067, K1095 at the HD2/TPR1 interface. F1092L may weaken these interactions and destabilize the interface. |
| E1136Q | TPR1 | TPR1 | Not clear. E1136 seems to interact with K1097, which in turn stacks with W1133 to stabilize the helix bundle of TPR1. E1136Q may form stacking interaction with W1133 directly and change its conformation, destabilizing the helix bundle of TPR1. |
| R1188P/Q | TPR1 | TPR1/NBD | R1188 is located in a loop that extends from TPR1 to interact with the NBD. R1188 is close to E803, D805 and E802 on NBD. R1188P may affect the conformation of the entire loop, while R1188Q may have weakened interaction. |
| D1190E/N | TPR1 | TPR1/NBD | Similar to R1188P, D1190E/N may affect the conformation of the loop that extends from TPR1 to NBD. |
| A1195V | TPR1 | TPR1/NBD? | A1195 is at the boundary of helix 1195-1214 and an N-terminal loop (which extends to NBD), and it is exposed on the surface. A1195V may change the loop-to-helix folding to hide the hydrophobic valine residue and affect the NBD/TPR1 loop binding. |
| Q1198H | TPR1 | TPR1 | Not clear. Q1198 seems to be involved in a network of hydrogen bonds with neighboring residues across several helices of TPR1. Several ordered water molecules are also involved in this interaction network. Q1198H may break this network. It may also repel the nearby K1295 and K1128. |
| Y1274C | TPR1 | TPR1/TPR2 | Y1274 is among a series of stacking residues that stabilize the TPR1-TPR2 interface, including P1280, K1475, Y1274, F1270, Y1197, Q1167. |
| V1276I | TPR1 | TPR1/TPR2 | V1276 is at the TPR1-TPR2 interface, which is stabilized by several pairs of interacting residues. The slightly bulkier V1276I residue may change the spacing of the two neighboring helices and affect the other interactions. |
| P1280L | TPR1 | TPR1/TPR2 | Similar to Y1274C. P1280 is among a series of stacking residues that stabilize the TPR1-TPR2 interface, including P1280, K1475, Y1274, F1270, Y1197, Q1167. |
| Q1286K | TPR1 | TPR1 | Not clear. Q1286 is located at the boundary of loop 1277-1286 and helix 1287-1309. Q1286K may affect the local folding. |
| <u>R1293Q/W</u>[a] | TPR1 | TPR1/TPR2 | R1293 forms a prominent stacking interaction with F1275, which in turn interacts with several other residues. Together, these interactions may have determined the relative positioning of helix 1287-1309 and helix 1261-1277 in an X shape. The loop connecting these two helices, 1278-1286, is the major contact point between TPR1 and TPR2 via two short beta strands. |

| | | | |
|---|---|---|---|
| V1329E | TPR2 | TPR2 | V1329 is embedded between two helices in TPR2. V1329E may affect the stability of TPR2 helix bundle. |
| K1353E | TPR2 | TPR1/TPR2 | K1353 is at the turning point of helix 1344-1352 and helix 1354-1374 in TPR2. It has a network of interactions with Y1350 (stacking), E1349 (salt-bridge), and a water molecule (hydrogen bond). E1349 is in turn interacting with R1293 across the TPR1-TPR2 interface. So K1353E may affect the TPR1-TPR2 interface by breaking the network of interactions. |
| N1387H | TPR2 | TPR1/TPR2 | N1387 stabilizes the conformation of loop 1341-1344 via hydrogen bonds with its mainchain atoms and water molecules. Loop 1341-1344 in turn contains several interactions with TPR1, such as D1342-K1265 and K1343-D1269. |
| S1463P | TPR2 | TPR2 | S1463 is in an α-helix of TPR2. S1463P may change the α-helical folding and destabilize TPR2. |
| I1478T | TPR2 | TPR1/TPR2/OB | I1478 is located in the middle of two short beta-strands--K1475-P1477 that augments with beta-strand L1278-P1280 in TPR1, and A1479-F1482 that augments with the beta-barrel of the OB fold. The sidechain of I1478 has stacking interactions with sidechains of K1279 and Y1480, as well as the mainchain peptide bond between L1278 and K1279. I1478T may break up this intricate balance and destabilize this multi-domain interface. |
| A1479S | TPR2 | TPR2/OB | A1479 has hydrophobic interactions with F1482 (on TPR2), Y1584 (on OB) and L1582 (on OB). A1479S may destabilize this hydrophobic TPR2-OB interface. |
| K1487R | TPR2 | local conformation | Not clear. K1487 is exposed on the surface of TPR2 and located in a loop. |
| S1514N | TPR2 | local conformation | Not clear. S1514 is located at the tip of a loop structure. |
| R1529H | OB | OB | R1529 may strengthen the OB fold by bundling the three neighboring beta-strands. It has stacking interaction with Y1573, hydrogen bond with S1571, hydrogen bond with the mainchain carbonyl of D1585, and salt bridge with the sidechain of D1585. |
| R1533Q | OB | OB/NBD, OB/SIR2 | Not clear. R1533Q may somehow promote the folding of the neighboring loop 1562-1569 into the OB/SIR2 interacting mode, which is also the OB/NBD disengaged mode. |
| L1539I | OB | OB/NBD, OB/SIR2 | Not clear. May be similar to R1533Q--promote the folding of the neighboring loop 1562-1569 into the OB/SIR2 interacting mode, which is also the OB/NBD disengaged mode. |

| | | | |
|---|---|---|---|
| G1564S | OB | OB/NBD, OB/SIR2 | The backbone of loop 1562-1569 has a 90-degree kink at G1564. The stacking interaction between the S1563-G1564 peptide bond and the sidechain of Y825 in NBD is one of the major OB/NBD interactions. G1564S may make the backbone more rigid and less preferrable to form the OB/NBD interaction. Or the sidechain of G1564S may form extra hydrogen bond at the OB/SIR2 interface. |
| I1567M | OB | OB/NBD, OB/SIR2 | Not clear. May destabilize the OB/NBD interaction mode of the OB loop 1562-1569. I1567 has hydrophobic interaction with F465 and Y505 of NBD in the OB/SIR2 interface. |

[a] Underlined mutations denote recurrent GoF variants identified in more than four independent cases.

**Supplemental file 1. (separate file)**

**Structure-based sequence alignment of representative mammalian SAMD9/9L proteins.**

The sequence alignment was generated using Aliview, and the figure was prepared with ESPript. Secondary structure elements of the human SAMD9 cryo-EM structure and corresponding residue numbers are shown above the alignment. The line below the residues denotes domain boundaries and is color-coded according to Fig. 1B. Black sphere symbols within the line indicate positions mutated in this study.

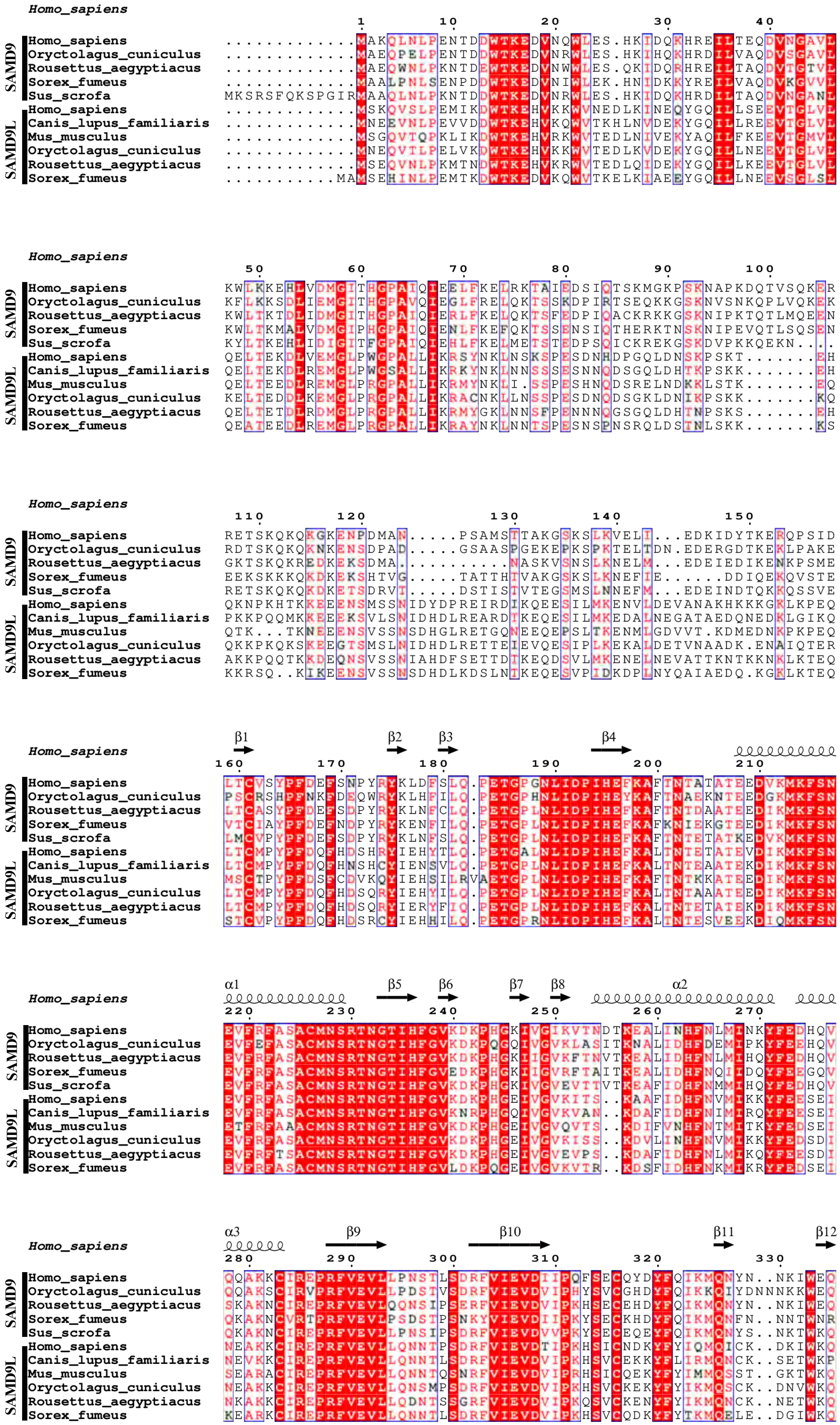

Homo_sapiens
1 10 20 30 40
SAMD9
Homo_sapiens .............MAKQLNLPENTDDWTKEDVNQWLES.HKIDQKHREILTEQDVNGAVL
Oryctolagus_cuniculus .............MAEQPELPENTDDWTKEDVNRWLES.HKIHQKHRDILVAQDVSGAVL
Rousettus_aegyptiacus .............MAEQWNLPKNTDEWTKEDVNWWLES.QKIDQRHREILTAQDVTGTVL
Sorex_fumeus .............MAALPNLSENPDDWTKEDVNIWLEK.HNIDKKYREILVAQDVKGVVL
Sus_scrofa MKSRSFQKSPGIRMAAQLNLPKNTDDWTKEDVNRWLES.HKIDQKHRDILTAQDVNGANL
SAMD9L
Homo_sapiens .............MSKQVSLPEMIKDWTKEHVKKWVNEDLKINEQYGQILSEEVTGLVL
Canis_lupus_familiaris .............MNEEVNLPEVVDDWTKEHVKQWVTKHLNVDEKYGQILREEVTGLVL
Mus_musculus .............MSGQVTQPKLIKDWTKEHVRKWVTEDLNIVEKYAQILFKEEVTGMVL
Oryctolagus_cuniculus .............MNEQVTLPELVKDWTKEHVKKWVTEDLKVDEKYGQILNEEVTGLVL
Rousettus_aegyptiacus .............MSEQVNLPKMTNDWTKEHVKRWVTEDLQIDEKYGQILKEEVTGLVL
Sorex_fumeus ...........MAMSEHINLPEMTKDWTKEDVKQWVTKELKIAEEYGQILNEEVSGLSL

Homo_sapiens
50 60 70 80 90 100
SAMD9
Homo_sapiens KWLKKEHLVDMGITHGPAIQIEELFKELRKTAIEDSIQTSKMGKPSKNAPKDQTVSQKER
Oryctolagus_cuniculus KFLKKSDLIEMGITHGPAVQIEGLFRELQKTSSKDPIRTSEQKKGSKNVSNKQPLVQKEK
Rousettus_aegyptiacus KWLTKTDLIDMGITHGPAIQIERLFKELQKTSFEDPIQACKRKKGSKNIPKTQTLMQEEN
Sorex_fumeus KWLTKMALVDMGIPHGPAIQIENLFKEFQKTSSENSIQTHERKTNSKNIPEVQTLSQSEN
Sus_scrofa KYLTKEHLIDIGITFGPAIQIEHLFKELMETSTEDPSQICKREKGSKDVPKKQEKN....
SAMD9L
Homo_sapiens QELTEKDLVEMGLPWGPALLIKRSYNKLNSKSPESDNHDPGQLDNSKPSKT.......EH
Canis_lupus_familiaris QELTEKDLREMGLPWGSALLIKRKYNKLNNSSSESNNQDSGQLDHTKPSKK.......EH
Mus_musculus QELTEEDLREMGLPRGPALLIKRMYNKLI.SSPESHNQDSRELNDKKLSTK.......EQ
Oryctolagus_cuniculus KELTEKDLKEMGLPRGPALLIKRACNKLLNSSPESDNQDSGKLDNIKPSKK.......KQ
Rousettus_aegyptiacus QELTETDLRDMGLPRGPALLIKRMYGKLNNSFPENNNQGSGQLDHTNPSKS.......EH
Sorex_fumeus QEATEEDLREMGLPRGPALLIKRAYNKLNNTSPESNSPNSRQLDSTNLSKK.......KS

Homo_sapiens
110 120 130 140 150
SAMD9
Homo_sapiens RETSKQKQKGKENPDMAN.....PSAMSTTAKGSKSLKVELI...EDKIDYTKERQPSID
Oryctolagus_cuniculus RDTSKQKQKNKENSDPAD.....GSAASPGEKEPKSPKTELTDN.EDERGDTKEKLPAKE
Rousettus_aegyptiacus GKTSKQKREDKEKSDMA...........NASKVSNSLKNELM...EDEIEDIKENKPSME
Sorex_fumeus EEKSKKKQKDKEKSHTVG.....TATTHTVAKGSKSLKNEFIE......DDIQEKQVSTE
Sus_scrofa RETSKQKQKDKETSDRVT.....DSTISTVTEGSMSLNNEFM...EDEINDTQKKQSSVE
SAMD9L
Homo_sapiens QKNPKHTKKEEENSMSSNIDYDPREIRDIKQEESILMKENVLDEVANAKHKKKGKLKPEQ
Canis_lupus_familiaris PKKPQQMKKEEEKSVLSNIDHDLREARDTKEQESILMKEDALNEGATAEDQNEDKLGIKQ
Mus_musculus QTK...TKNEEENSVSSNSDHGLRETGQNEEQEPSLTKENMLGDVVT.KDMEDNKPKPEQ
Oryctolagus_cuniculus QKKPKQKSKEEGTSMSLNIDHDLRETTEIEVQESIPLKEKALDETVNAADK.ENAIQTER
Rousettus_aegyptiacus AKKPQQTKKDEQNSVSSNIAHDFSETTDTKEQDSVLMKENELNEVATTKNTKKNKLKTEQ
Sorex_fumeus KKRSQ..KIKEENSVSSNSDHDLKDSLNTKEQESVPIDKDPLNYQAIAEDQ.KGKLKTEQ

β1 β2 β3 β4
Homo_sapiens
160 170 180 190 200 210
SAMD9
Homo_sapiens LTCVSYPFDEFSNPYRYKLDFSLQ.PETGPGNLIDPIHEFKAFTNTATATEEDVKMKFSN
Oryctolagus_cuniculus PSCRSHPFNKFDEQWRYKLHFILQ.PETGPHNLIDPIHEYKAFTNAEKNTEEDGKMKFSN
Rousettus_aegyptiacus LTCASYPFDEFSDPYRYKLNFCLQ.PETGPLNLIDPIHEFKAFTNTDAATEEDIKMKFSN
Sorex_fumeus VTCIAYPFDEFNDPYRYKENFILQ.PETGPLNLIDPIHEFKAFKNIEKGTEEDIKMKFSN
Sus_scrofa LMCVPYPFDEFSDPYRYKLNFSLQ.PETGPLNLIDPIHEFKAFTNTETATKEDVKMKFSN
SAMD9L
Homo_sapiens LTCMPYPFDQFHDSHRYIEHYTLQ.PETGALNLIDPIHEFKALTNTETATEVDIKMKFSN
Canis_lupus_familiaris LTCMPYPFDQFHNSHCYIENSVLQ.PETGPLNLIDPIHEFKALTNTEAATEKDIKMKFSN
Mus_musculus MSCTPYPFDSFCDVKQYIEHSILRVAETGPLNLIDPIHEFKAFTNTKKATEEDIKMKFSN
Oryctolagus_cuniculus LTCMPYPFDQFHDSQRYIEHYILQ.PETGPLNLIDPIHEFKALTNTAAATEEDIKMKFSN
Rousettus_aegyptiacus LTCMPYPFDQFHDSQRYIERYFIQ.PETGPLNLIDPIHEFKALTNTETATEKDIKMKFSN
Sorex_fumeus STCVPYPFDQFHDSRCYIEHHILQ.PETGPRNLIDPIHEFKALTNTESVEEKDIQMKFSN

α1 β5 β6 β7 β8 α2
Homo_sapiens
220 230 240 250 260 270
SAMD9
Homo_sapiens EVFRFASACMNSRTNGTIHFGVKDKPHGKIVGIKVTNDTKEALINHFNLMINKYFEDHQV
Oryctolagus_cuniculus EVFEFASACMNSRTNGTIHFGVKDKPQGQIVGVKLASITKNALIDHFDEMIPKYFEEHQV
Rousettus_aegyptiacus EVFRFASACMNSRTNGTIHFGVKDKPHGKIIGVKFTNVTKEALIDHFNLMIHQYFEDHQV
Sorex_fumeus EVFRFASACMNSRTNGTIHFGVEDKPHGKIIGVRFTAITKEALIDHFNQIIDQYFEEGQV
Sus_scrofa EVFRFASACMNSRTNGTIHFGVKDKPHGKIVGVEVTTVTKEAFIDHFNMMIHQYFEDHQV
SAMD9L
Homo_sapiens EVFRFASACMNSRTNGTIHFGVKDKPHGEIVGVKITS..KAAFIDHFNVMIKKYFEESEI
Canis_lupus_familiaris EVFRFASACMNSRTNGTIHFGVKNRPHGQIVGVKVAN..KDAFIDHFNIMIRQYFEESEI
Mus_musculus ETFRFAAACMNSRTNGTIHFGVKDKPHGEIVGVQVTS..KDIFVNHFNTMITKYFEDSEI
Oryctolagus_cuniculus EVFRFASACMNSRTNGTIHFGVKDKPHGEIVGVKISS..KDVLINHFNVMIKKYFEDSDI
Rousettus_aegyptiacus EVFRFTSACMNSRTNGTIHFGVKDKPHGEIVGVEVPS..KDAFIDHFNLMIKQYFEESDI
Sorex_fumeus EVFRFASACMNSRTNGTIHFGVLDKPQGEIVGVKVTR..KDSFIDHFNKMIKRYFEDSEI

α3 β9 β10 β11 β12
Homo_sapiens
280 290 300 310 320 330
SAMD9
Homo_sapiens QQAKKCIREPRFVEVLLPNSTLSDRFVIEVDIIPQFSECQYDYFQIKMQNYN..NKIWEQ
Oryctolagus_cuniculus QQAKSCIRVPRFVEVLLPDSTVSDRFVIEVDVIPHYSVCGHDYFQIKKQIYDNNNKKWEQ
Rousettus_aegyptiacus SKAKNCIREPRFVEVLQQNSIPSERFVIEVDVIPKHSECEHDYFQIKMQNYN..NKIWEQ
Sorex_fumeus QKAKNCVRTPRFVEVLPSDSTPSNKYVIEVDIIPKYSECKHDYFQIKMQNFN..NNTWNR
Sus_scrofa QKAKNCIREPRFVEVLLPNSIPSDRFVIEVDVVPKYSECEQEYFQIKMQNYS..NKTWKQ
SAMD9L
Homo_sapiens NEAKKCIREPRFVEVLLQNNTPSDRFVIEVDTIPKHSICNDKYFYIQMQICK..DKIWKQ
Canis_lupus_familiaris NEVKKCIREPRFVEVLLQNNTLSDRFVIEVDVIPKHSVCEKKYFLIRMQNCK..SETWKP
Mus_musculus SEARACIREPRFVEVLLQNNTQSNRFVIEVDVIPRHSICQEKYFYIMMQSST..GKTWKQ
Oryctolagus_cuniculus NEAKKCIREPRFVEVLLQNSMPSDRFVIEVDVIPKHSVCKEKYFYIKMQSCK..DNVWKQ
Rousettus_aegyptiacus NKAKKCIREPRFVEVLLQDNTSSGRFVIEVDVIPKQSVCKENYFYIKMQNCK..NETWKQ
Sorex_fumeus KEARKCIREPRFVEVLLQNNTLSDRFVIEVDVIPKHSVCQDKYFYTKMQELE..DGIWKQ

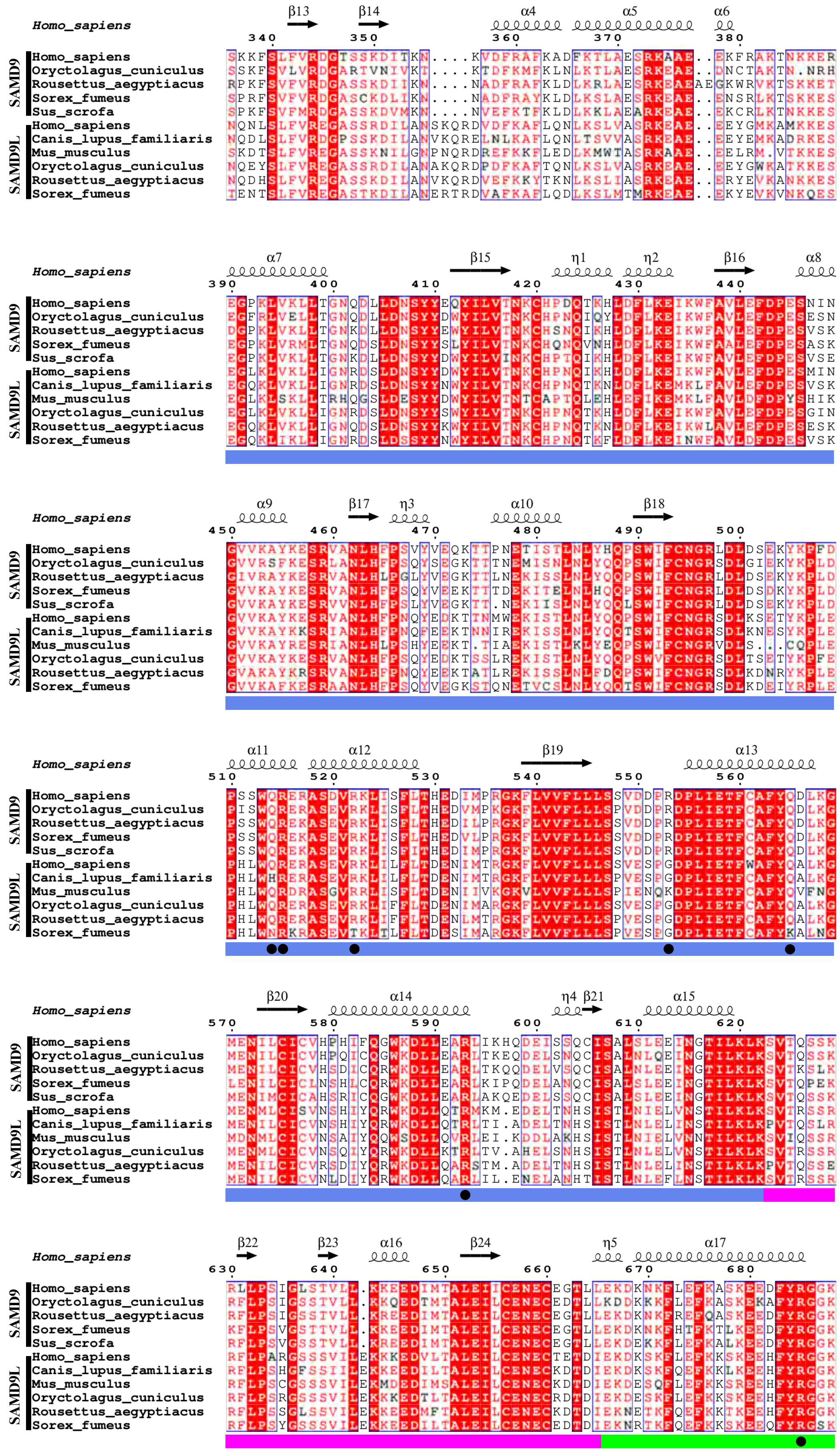
Homo_sapiens
SAMD9
Homo_sapiens
Oryctolagus_cuniculus
Rousettus_aegyptiacus
Sorex_fumeus
Sus_scrofa
SAMD9L
Homo_sapiens
Canis_lupus_familiaris
Mus_musculus
Oryctolagus_cuniculus
Rousettus_aegyptiacus
Sorex_fumeus
β13 β14 α4 α5 α6
α7 β15 η1 η2 β16 α8
α9 β17 η3 α10 β18
α11 α12 β19 α13
β20 α14 η4 β21 α15
β22 β23 α16 β24 η5 α17

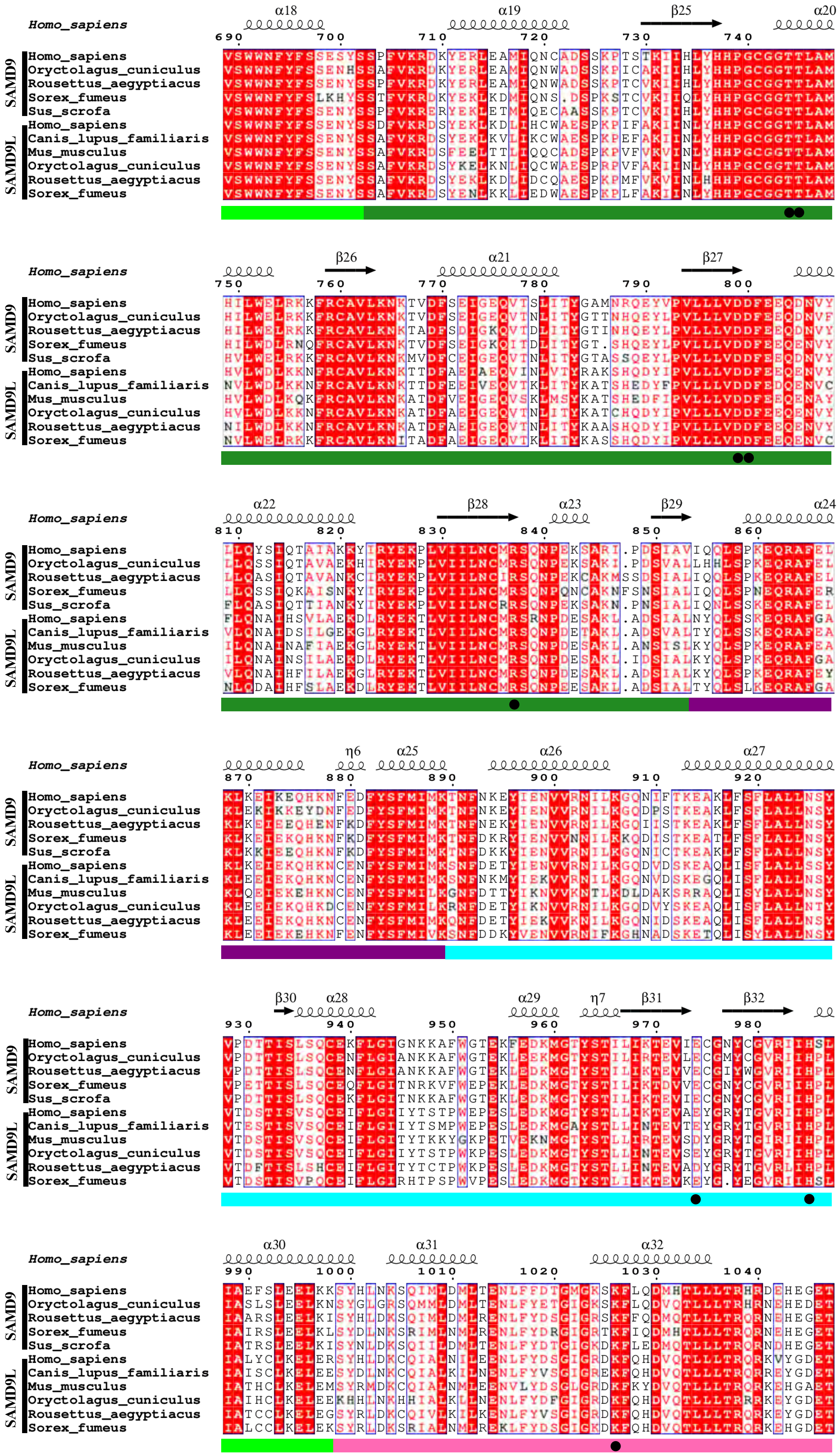
Homo_sapiens
SAMD9
Homo_sapiens
Oryctolagus_cuniculus
Rousettus_aegyptiacus
Sorex_fumeus
Sus_scrofa
SAMD9L
Homo_sapiens
Canis_lupus_familiaris
Mus_musculus
Oryctolagus_cuniculus
Rousettus_aegyptiacus
Sorex_fumeus
α18 α19 β25 α20
β26 α21 β27
α22 β28 α23 β29 α24
η6 α25 α26 α27
β30 α28 α29 η7 β31 β32
α30 α31 α32

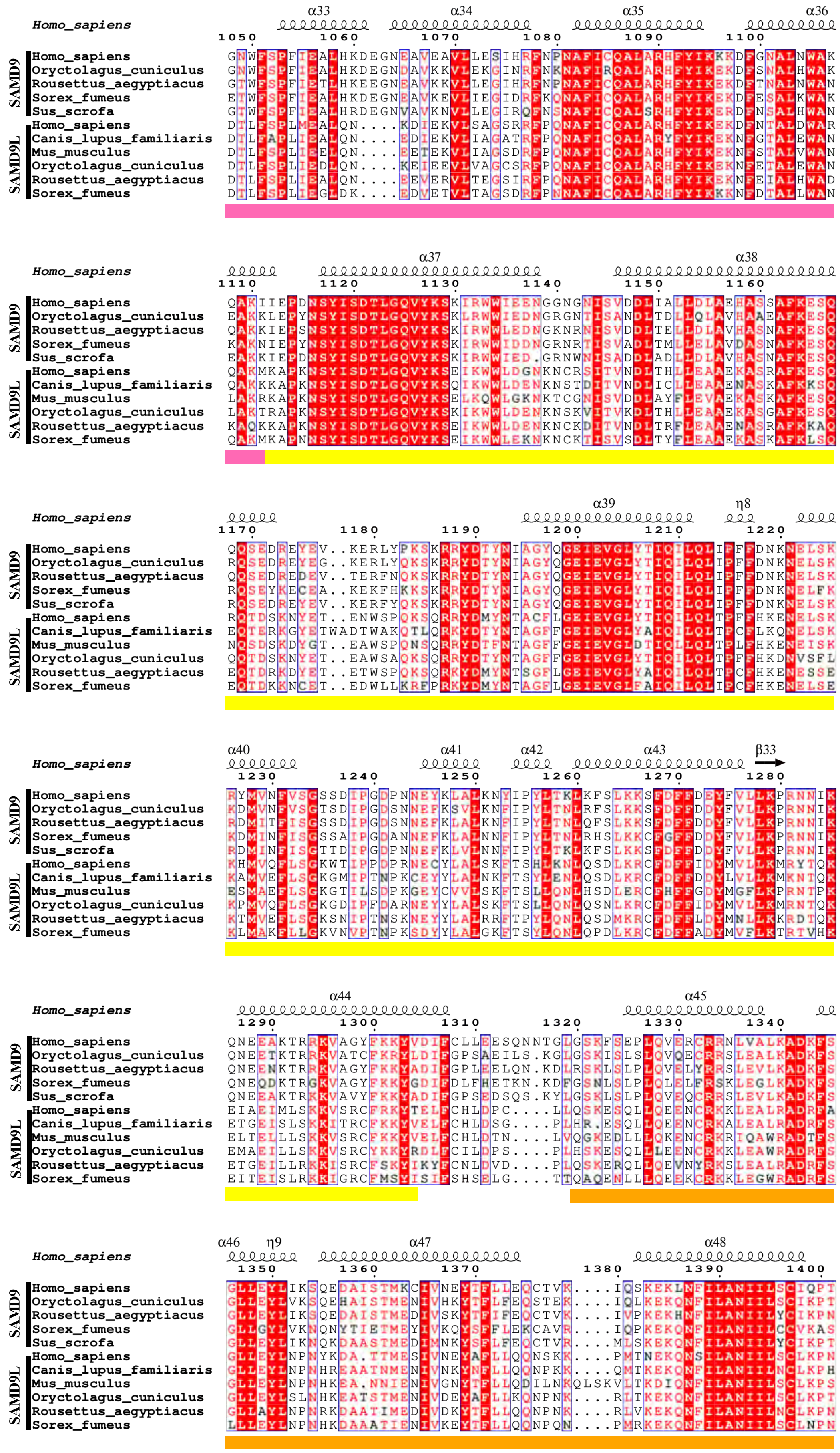
Homo_sapiens
SAMD9
SAMD9L
Homo_sapiens
Oryctolagus_cuniculus
Rousettus_aegyptiacus
Sorex_fumeus
Sus_scrofa
Canis_lupus_familiaris
Mus_musculus

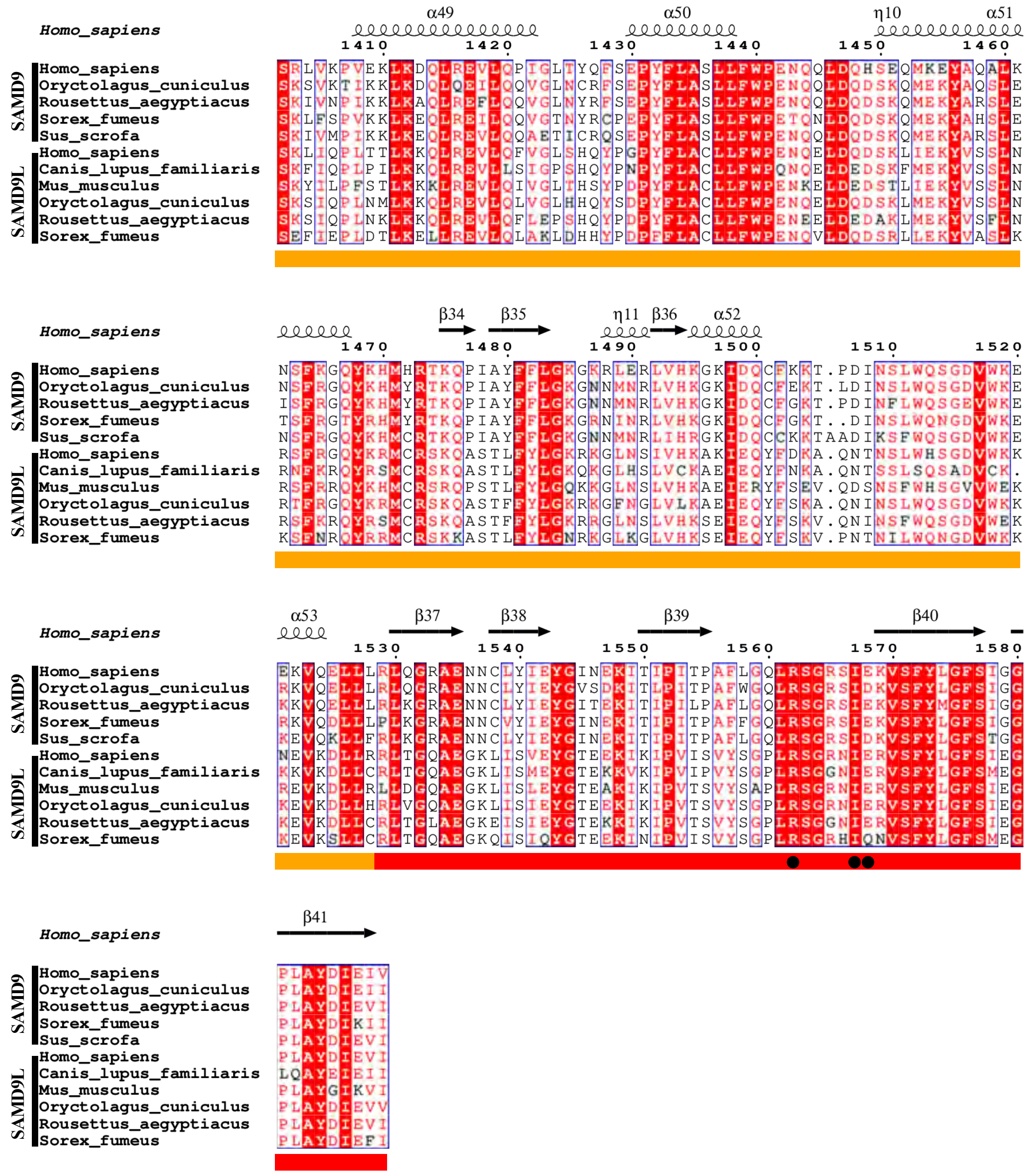
Homo_sapiens
α49 α50 η10 α51
1410 1420 1430 1440 1450 1460
β34 β35 η11 β36 α52
1470 1480 1490 1500 1510 1520
α53 β37 β38 β39 β40
1530 1540 1550 1560 1570 1580
β41
SAMD9
Homo_sapiens
Oryctolagus_cuniculus
Rousettus_aegyptiacus
Sorex_fumeus
Sus_scrofa
SAMD9L
Homo_sapiens
Canis_lupus_familiaris
Mus_musculus
Oryctolagus_cuniculus
Rousettus_aegyptiacus
Sorex_fumeus
